# Tuning movement for sensing in an uncertain world

**DOI:** 10.1101/826305

**Authors:** Chen Chen, Todd D. Murphey, Malcolm A. MacIver

## Abstract

While animals track or search for targets, sensory organs make small unexplained movements on top of the primary task-related motions. While multiple theories for these movements exist—in that they support infotaxis, gain adaptation, spectral whitening, and high-pass filtering—predicted trajectories show poor fit to measured trajectories. We propose a new theory for these movements called energy-constrained proportional betting, where the probability of moving to a location is proportional to an expectation of how informative it will be balanced against the movement’s predicted energetic cost. Trajectories generated in this way show good agreement with measured target tracking trajectories of electric fish. Similarly good agreement was found across three published datasets on visual and olfactory tracking tasks in insects and mammals. Our theory unifies the metabolic cost of motion with information theory. It predicts sense organ movements in animals and can prescribe sensor motion for robots to enhance performance.

## Introduction

Movement can be used to obtain information that is not uniformly distributed in the environment. Because movement is energetically costly, there is likely a balance between the benefits of increased sensory information and energetic costs for obtaining that information. We have developed a theory that unifies these two dimensions of information acquisition and can be applied across sensory modalities and species. This theory, energy-constrained proportional betting, predicts the small and seemingly extraneous movements that sensory organs or animals undergo as they near or track a target of interest (***Martin, 1965**; **Basil et al., 2000**; **Ferner and Weissburg, 2005**; **Webb et al., 2004**; **Willis and Avondet, 2005**; **Porter et al., 2007**; **Louis et al., 2008**; **Duistermars et al., 2009**; **Yovel et al., 2010**; **Khan et al., 2012**; **Stamper et al., 2012**; **Catania, 2013**; **Sponberg et al., 2015**; **Lockey and Willis, 2015**; **Rucci and Victor, 2015**; **Stockl et al., 2017***) (see Figure 1 — figure supplement 1). These movements appear unrelated to the movements that are necessary to achieve the task at hand. For example, weakly electric fish will track and stay near a moving refuge, but in addition to the large motions needed to stay near the refuge there are small wholebody oscillations, an electrosensory analog to microsaccades (***Stamper et al., 2012***) (see Figure 1 — figure supplement 1). Similarly, in behaviors where animals sample discretely over time, animals vary their sampling frequency or the location at which samples are taken, as observed in bats, rats, beaked whales, humans, and pulse electric fish (***Yovel et al., 2010**; **Mitchinson et al., 2007**; **Hartmann, 2001**; **Kothari et al., 2018**; **Caputi et al., 2003**; **Pluta and Kawasaki, 2008**; **Nelson and MacIver, 2006**; **Schnitzler et al., 2003**; **Madsen et al., 2005**; **Yang et al., 2016**; **Hoppe and Rothkopf, 2019**).*

There have been several theories proposed to account for these sensing movements including information maximization or infotaxis (***Najemnik and Geisler, 2005**; **Vergassola et al., 2007**; **Yovel et al., 2010**; **Calhoun et al., 2014**; **Álvarez-Salvado et al., 2018**; **Yang et al., 2016**),* gain adaptation (***Stockl et al., 2017**; **Biswas et al., 2018**),* spectral whitening (***Rucci and Victor, 2015**),* and high-pass filtering (***Stamper et al., 2012***). However, most existing theories are underspecified in that they do not attempt to provide a complete control framework, and are therefore incapable of generating realistic trajectories to facilitate direct behavioral validation of the model (see Discussion). We show that an implementation of energy-constrained proportional betting generates trajectories with good agreement to measured behavior.

An important quantity for implementing energy-constrained proportional betting is the expected information density (EID). To understand it, imagine that you are trying to locate a wi-fi router hidden directly under a long narrow counter along which you are moving your laptop by way of the signal strength reported by your laptop. The router emits a signal and one can model how strong that signal will be as a function of the location of the laptop (the sensor) and the router (the object). This model relating the sensor reading to the object location is called the observation model. In this case, it may be captured by a set of (position along counter, expected reading) values—called a 1-D observation model. All the observation models used for generating the results shown in this study are similarly also 1-D. As one moves along the counter, the signal strength varies according to the 1-D observation model, with some noise degrading the signal strength measurement. To get the best signal, one would likely move directly to the current best estimate of the router location. But to get a better estimate of router location, moving to a counter position that maximizes the information about the router location makes sense—and is generally not going to be the same as moving to the estimated router location. Instead, places where the signal strength is most sensitive to changes in location provide the most evidence about router location. The information density will be highest at those places; since those locations are not known, one has to compute the expected information density based on the observation model conditioned on where the router is expected to be based on current evidence. For a graphical explanation of the EID, see Video 1.

Let’s assume that we have a way to compute the EID for a given target—a 1-D quantity for a target along a line, a 2-D quantity for a target on a surface as shown in Figure 1A, and a 3-D quantity for a target in space. One possible way to generate target-related behavior is to monitor a single peak in the EID to greedily maximize the information gain (Figure 1 B). A method called infotaxis (***Vergassola et al., 2007***) similarly generates trajectories that maximize expected information by commanding motion toward a peak of the EID. However, expected information maximization leads to problems when there is a high level of uncertainty, as is frequently the case in naturalistic conditions.In such conditions, the concept of distractors becomes relevant. A distractor represents either a real physical object which appears similar to the true target, or a transient object-like appearance caused by noise (we will term these fictive distractors to avoid conflating these with physical distractors). Figure 1 B shows the behavior of an expected information maximizing solution in the presence of a distractor. Because the gradient of increased expected information leads to the distractor, the sensor is commanded to go straight to it.

**Figure 1.**
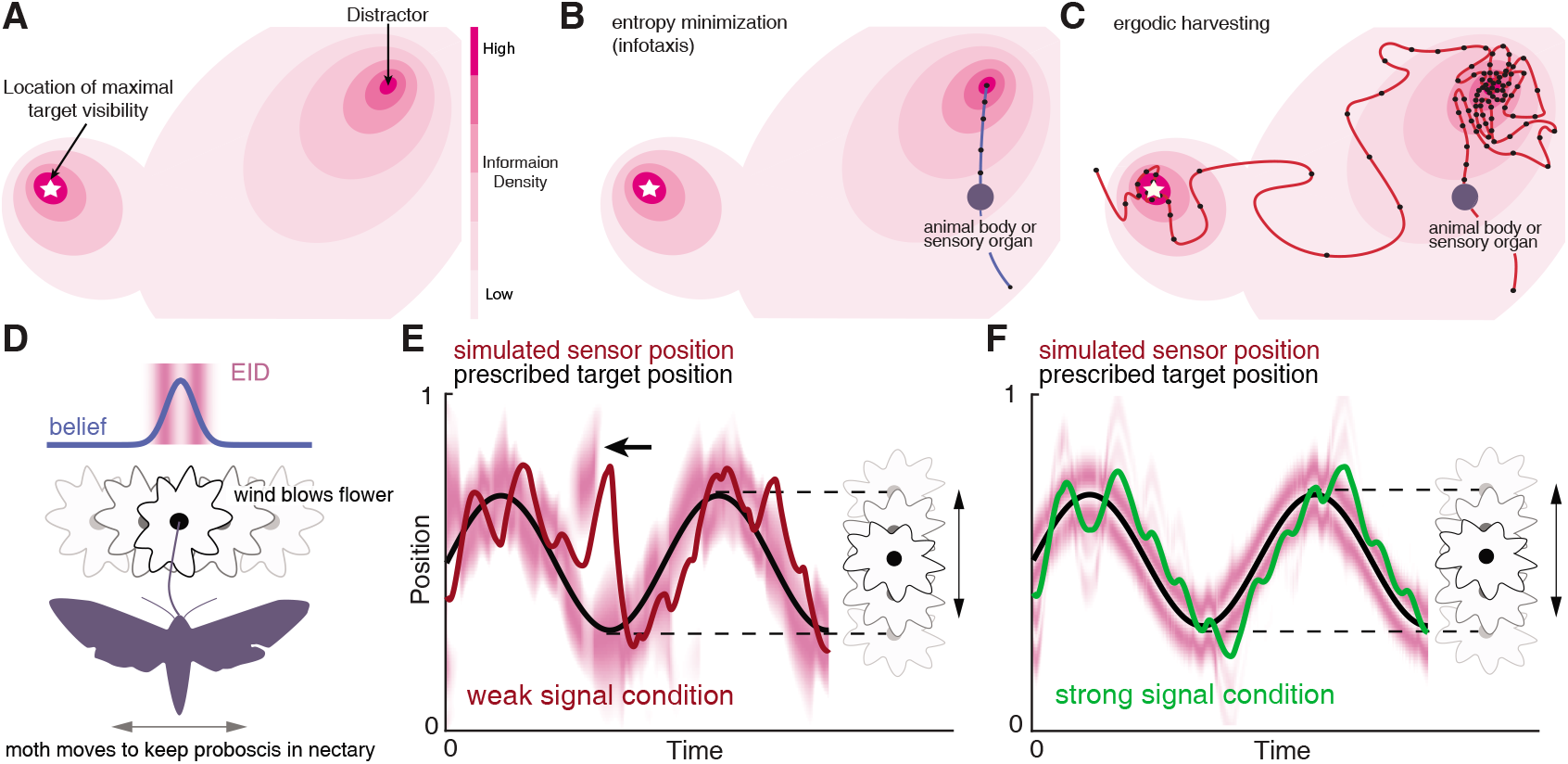
Illustration of a 2-D expected information density, information maximization and energy-constrained proportional betting. (**A**) The heat map represents the expected information density. Because the peak expected information is typically not at the same location as the object, we illustrate the target peak as the point of maximum target visibility. (**B**) Trajectories generated by information maximization (entropy minimization) locally maximizes the expected information at every step, which here commands a path straight to the nearby distractor. In contrast, energy-constrained proportional betting (**C**) samples the EID proportionate to its density but balanced by the cost of movement. The natural tradeoff between exploration and exploitation that emerges leads to localization of the target and rejection of the distractor (adapted from Figure 3 of ***Miller et al. 2016***). Black dots along the trajectory indicate samples at fixed time intervals (longer distances between dots indicate higher speed). (**D-F**) An illustration of an animal tracking an object constrained to move in a line, in this case a hypothetical moth following a flower swaying in a breeze in a manner approximated by a 1-D sinusoid—a natural behavior reported in ***Sponberg et al. (2015); Stockl et al. (2017)***. We simulate the tracking of the flower using energy-constrained proportional betting. (**D**) In the top panel, we show an idealization of the moth’s belief (blue line Gaussian distribution above the moth) about the flower’s location when the flower reaches the center point. Higher values in the *y* direction represents higher confidence of the target at the given *x* location. The corresponding EID is overlaid in magenta;a darker color indicates higher expected information should the animal take a new sensory measurement in the corresponding location. Note the bimodal structure of this highly idealized EID, with identical maximal information peaks on both sides of the Gaussian belief. Intuitively, the expected information is higher along the high slope region of the belief because in this region, small changes in location (x axis direction in the upper inset) will cause large changes in the received sensory signal and hence carry more information about position. (**E**) Simulation of the moth’s position (red curve) while tracking a moving target (black curve) under a weak signal condition (Methods). The corresponding EID is overlaid in magenta. Note that even though trajectory segments are planned at 14 fixed-time intervals *T* (Methods) over the shown duration based on the EID at the start of those intervals, the EID is here plotted continuously for visualization purposes only. Energy-constrained proportional betting results in persistent activation of movement when the EID is relatively diffuse due to lack of information. Note the presence of a fictive distractor (marked by the black arrow) in the EID due to higher uncertainty in the sensory input as a result of the weaker signal. As seen by the digression below the black arrow, EIH responded to the distractor by making a detour away from the actual target position to gamble on the chance of acquiring more information, but does not get trapped by the distractor because of the proportional betting strategy in combination with the transient nature of fictive distractors. (**F**) Same as (E), but under strong signal conditions. Now the energy-constrained proportional betting trajectory samples both peaks of the bimodal EID with excursions away from these peaks, similar to measured moth behavior (***Stockl et al. 2017***, see Figure 1—fìgure supplement 1 “Moth”). The following figure supplements are available for figure 1: **Figure 1—fìgure supplement 1.** Whole-body or sensory organ small-amplitude motions are ubiquitous as animals track targets.

In contrast to the expected information maximizing solution, with energy-constrained proportional betting, sensory organs (or signal emitters in the case of animals like bats and electric fish) are moved to sample spatially distributed signals proportionate to the EID as shown in Figure 1C, constrained by the energetic cost of movement. The underlying sensory sampling strategy gambles on the chance of obtaining more information at a given location through carefully controlled sensor motion that balances two factors that typically push in opposite directions: 1) proportionally bet on the expected information gain (that is, take more samples by moving slowly in high EID regions, and fewer samples by moving more rapidly in low EID regions); and 2) minimize the energy expended for motion.

For this study we have quantified the expected informativeness of sensing locations by how much an observation at a location would reduce the Shannon-Weaver entropy (hereafter entropy) of the current estimate of the target’s location, as in infotaxis (***Vergassola et al., 2007**).* However, other measures of information such as Fisher Information can be used with near identical results— see ***Miller et al. 2016***. In our approach, the proximity of a given trajectory to perfect proportional betting is quantified by the ergodic metric. The ergodic metric provides a way of comparing a trajectory to a distribution (*e.g.,* the EID) by asking whether a trajectory over some time interval has the same spatial statistics as a given distribution (Methods, Appendix 3). Comparing a trajectory to a distribution is a novel capability of the ergodic metric (***Mathew and Mezic, 2011***) that is not shared by common methods of comparing two probability distributions (Appendix 2). Through optimizing a mathematical function that combines ergodicity with the energy of movement (see Algorithm 1) we obtain trajectories that bet on information balanced by the metabolic cost to move to informative locations in the space. With a perfectly ergodic trajectory (one with an ergodic measure of zero, only possible with infinite time and when the energy of movement is not considered), the distribution of expected information is perfectly encoded by the trajectory, or, equivalently, the trajectory does perfect proportional betting on the EID. We therefore call the associated algorithm ergodic information harvesting (hereafter EIH, modified from ***Miller et al. 2016***, see Methods). For a graphical explanation of EIH, see Video 2.

### Algorithm 1 Animal Tracking Simulation with Ergodic Information Harvesting

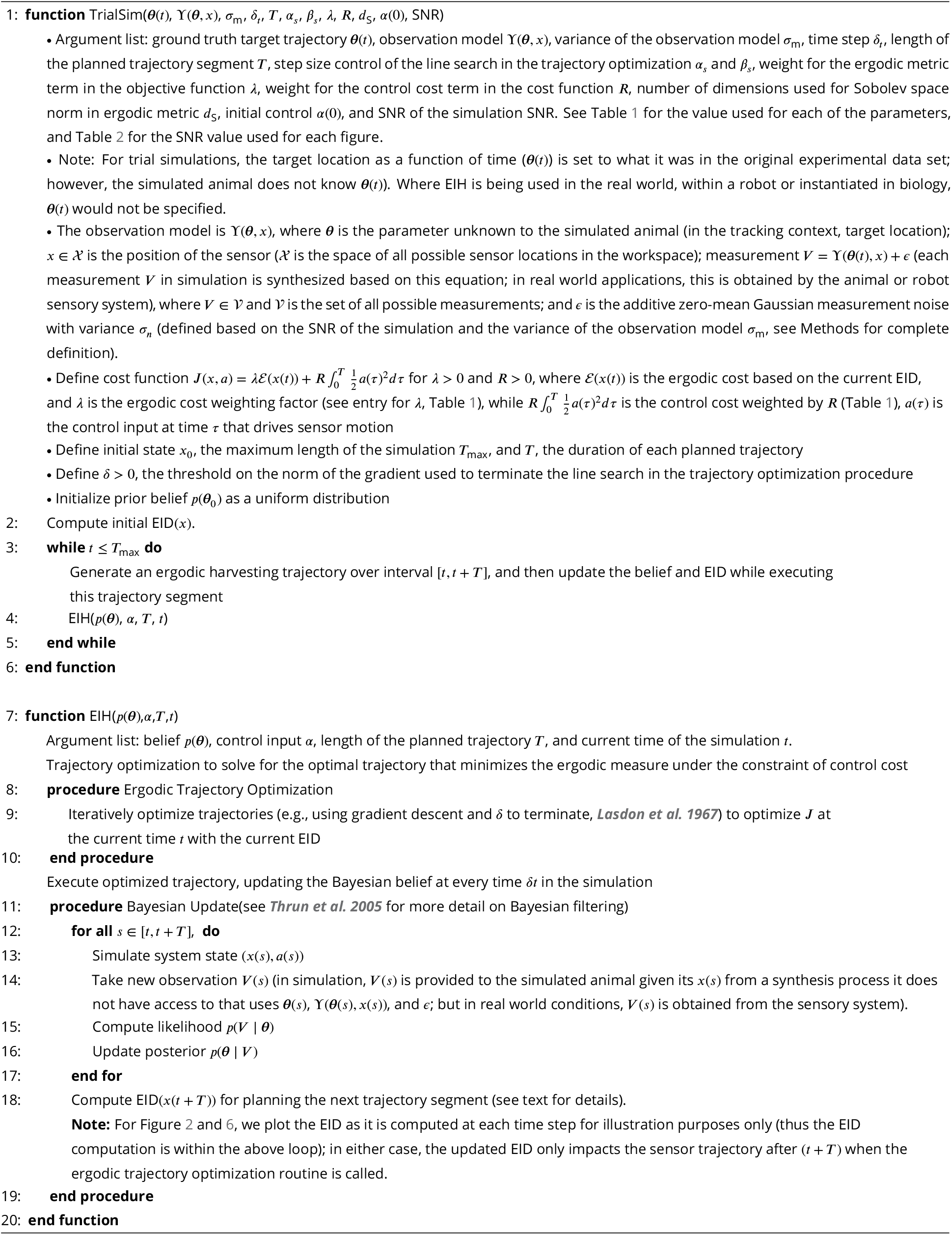

Figure 1E–F illustrates the emergence of oscillatory sensing-related motions as a simulated animal tracks an object moving back and forth along a line in a sinusoidal fashion using EIH. A key behavioral signature of EIH—increasing magnitude of sensing-related movements as signal weakens—is evident in this illustration, in contrast with the cessation of motion that occurs under very weak signals with information maximization (see Figure 6—figure supplement 4).

While prior studies have indicated that proportional betting is used at the cognitive decisionmaking level in primates (***Monosov et al., 2015**; **Gottlieb et al., 2014**),* our results suggest that an energy-constrained form of it occurs more broadly as an embodied component of information porcessing across a wide phylogenetic bracket. Below we will show evidence for this claim by comparing measured tracking trajectories to those simulated with ergodic information harvesting. Our core results use refuge tracking in weakly electric fish, but at the end we extend our results to three additional previously published datasets encompassing visual and olfactory tracking in insects and mammals.

## Results

First, we present a side-by-side comparison between the one-dimensional tracking trajectories generated by EIH and those we collected from South American gymnotid electric fish (glass knifefish *Eigenmannia virescens,* Valenciennes 1836) as they used electrosense to track a moving refuge in the dark (Figure 2A–D). Second, to examine how well EIH generalizes to other animals with different sensory modalities, we present similar comparisons between EIH and previously published behavioral datasets. These datasets were from blind eastern American moles (*Scalopus aquaticus,* Linnaeus 1758) finding an odor source (***Catania, 2013***); the American cockroach (*Periplaneta americana,* Linnaeus 1758) tracking an odor (***Lockey and Willis, 2015***); and the hummingbird hawkmoth (*Macroglossum stellatarum,* Linnaeus 1758) using vision and mechanosensory cues to track a swaying nectar source while feeding (***Stockl et al., 2017***). In all cases, animals were either tracking a moving target (electric fish and moth), or localizing a stationary target (mole and cockroach). Each of the live animal behavior datasets include experiments where the signal versus noise level of the dominant sensory modality driving the behavior was varied. For comparing the resulting trajectories against the predicted trajectory from EIH, this dominant sensory modality was selected for modeling.

**Figure 2.**
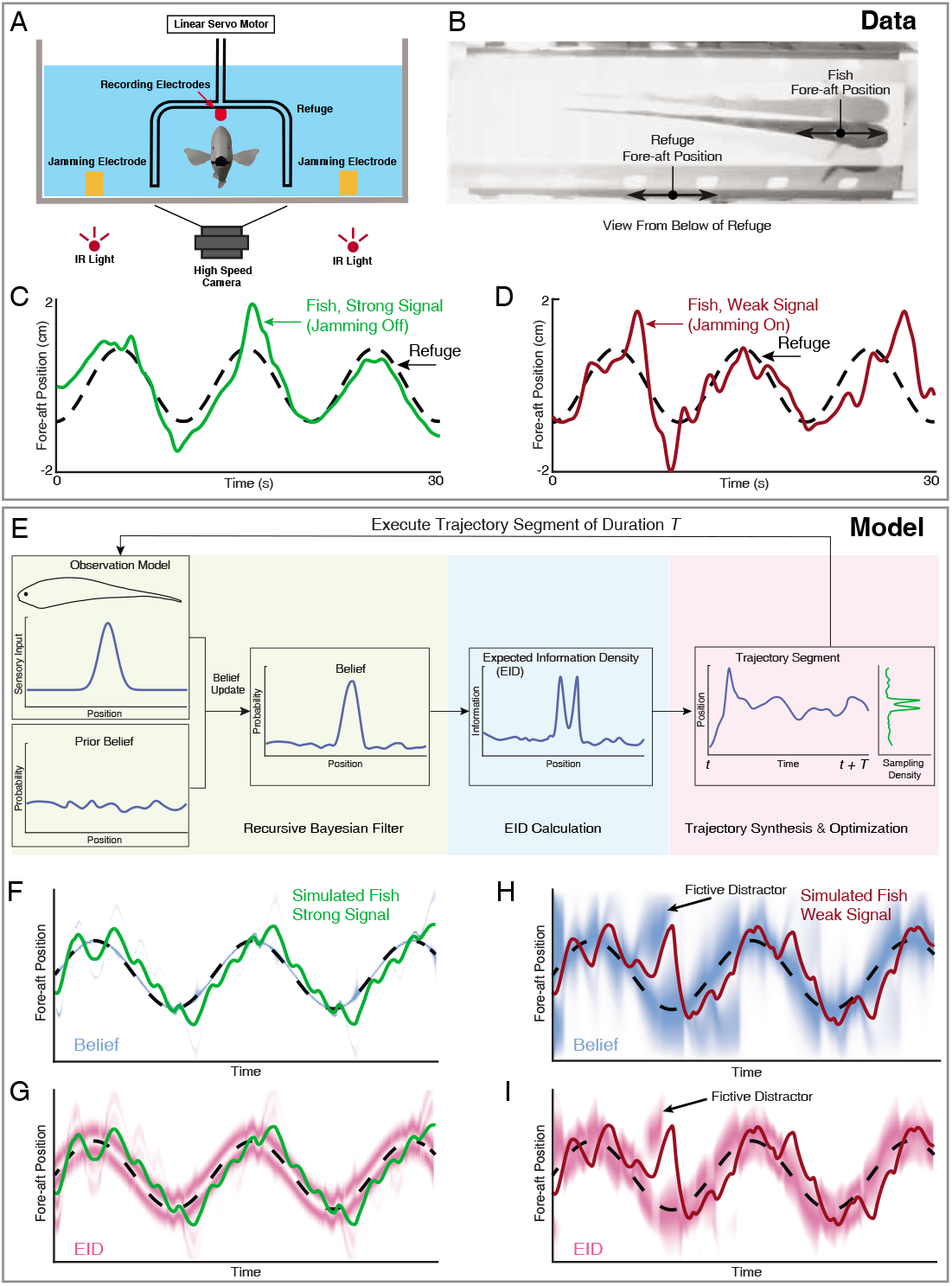
Longitudinal refuge tracking behavior in weakly electric fish and illustration of ergodic information harvesting. (**A**) Head-on view of experimental apparatus. A computer-controlled linear servo moves the refuge forward and backward along the longitudinal axis of the fish. Jamming electrodes are mounted to the side of the tank and recording electrodes are mounted to the ends of the refuge to generate and monitor the level of electrosensory noise (Methods). (**B**) An example frame of the captured video. (**C-D**) Fish tracking the refuge in the strong signal condition (no jamming applied), and in the weak signal condition (with jamming). Note that departures from the refuge position occur more often and with larger amplitude in the weak signal condition (***Stamper et al., 2012***). (**E**) Core components of the EIH algorithm, shown by colored blocks: a recursive Bayesian filter (green block), an expected information density (EID) calculation (blue block), and synthesis and optimization of a trajectory segment (pink block). The process starts with the simulated fish receiving new sensory input via its position and the observation model. Through the recursive Bayesian filter process the simulated fish updates its prior belief about the refuge’s true location (the belief is initially uniform since the location is unknown) and captures that increase in information in a posterior belief. It then computes the EID based on the posterior belief, explaining why the EID is bimodal when the variance in the estimate is low for the Gaussian observation model: Figure 1D and **G**. A trajectory segment is then computed that balances proportional betting on the EID (ergodicity) with the cost of movement. The resulting trajectory samples locations proportionate to the EID, as shown by the Sampling Density plot. After these three stages, the simulated fish executes the trajectory segment and returns to begin the process again. (**F, G**) Simulated fish behavior in the strong signal condition. The blue heatmap shows belief (top) of where the target is, where darker colors represent locations with high probability, and expected information density (bottom) is represented by the magenta heatmap where darker colors represent higher expected information. (**H, I**) As in **F, G**, but for the weak signal condition. As in the experimental observations, the departures from the refuge’s path are larger and more frequent in the weak signal condition. Note the presence of various color bands representing fictive distractors. The following figure supplements are available for figure 2: **Figure 2—figure supplement 1.** Dual target tracking simulation with EIH and infotaxis. **Figure 2—figure supplement 2.** Single target tracking simulation with EIH and infotaxis in the presence of a simulated physical distractor. **Figure 2—figure supplement 3.** Effect of jamming and how it varies with jamming intensity.

Each sensory system was modeled as a 1-D point-sensor with a Gaussian observation model— a deterministic map—that relates the sensory signal value to the variable that the animal is trying to estimate—here assumed to be the position of the target (see Methods). Sensor measurements were simulated by drawing values from the observation model given the sensor’s position relative to the target. We added normally distributed measurement noise—also described by a Gaussian function—with variance determined by a specified signal-to-noise ratio (SNR, Methods) to simulate the strong and weak signal conditions present in the live animal trials. These simulated measurements were used to update a probability distribution (often multi-peaked, such as shown in Figure 2H at fictive distractor) representing the simulated animal’s belief about the target’s likely location through a Bayesian update (***Thrun et al., 2005***) of the previous estimate (Methods). To generate a trajectory for sensory acquisition, at each planning update (Methods), the EIH algorithm takes the updated belief and calculates an EID to estimate information density as a function of location (Figure 2G & I, the result of the “EID Calculation” in Figure 2E). Then we generate an ergodic trajectory segment with respect to the EID to simulate the collection of more measurements. Throughout the Results, we show trajectory plots along with the EID heat map—similar to Figure 2G & I—to indicate the relationship between sensing-related movements and the EID as EIH carries out proportional betting with respect to the EID.

### Animals and EIH respond to weak signal conditions with increased exploratory movement

We first examined the weakly electric fish’s tracking behavior under strong and weak signal conditions (Figure 2A-D). Weakly electric fish engage in a behavior termed refuge tracking where they try to maintain their position inside a close-fitting open-ended enclosure—such as a plastic tube— even as that enclosure is translated forward and backward along the lengthwise axis of the fish (***Rose and Canfield, 1993***). Refuge tracking is a natural behavior within protective cover swayed by water flow, such as vegetation or root masses, during the fish’s inactive (diurnal) periods in the South American rivers in which they live (***Rose and Canfield, 1993***). Prior work has shown that as sensory input is degraded, these fish will engage in larger full-body excursions from the path taken by the robotically controlled refuge (***Stamper et al., 2012**; **Biswas et al., 2018**; **Rose and Canfield, 1993***). For the trials reported here, all in the dark under infrared illumination, we degraded electrosensory input through varying the intensity of an externally imposed electrical jamming stimulus (Methods) which has previously been shown to impair electrolocation performance (***Watanabe and Takeda, 1963**; **Bastian, 1987**; **Ramcharitar et al., 2005***). Two representative fish behavior tracking trials are shown in Figure 2C-D. The weak signal condition (Figure 2D) resulted in more body movement during tracking.

In Figure 2F-I, we show the corresponding EIH output when EIH is given the same target trajectory, under simulated strong and weak signal conditions. In these simulations, although the simulated fish experiment has the target location provided, the EIH algorithm does not know this location and is only given simulated measurements (see Algorithm 1). The progression of the belief distribution over time is shown in Figure 2F & G for strong and weak signal conditions, respectively. Immediately below the belief plot, we show the same trajectory with the corresponding EID visualized as the magenta overlay.

In order to quantify the increase in movement during tracking, we defined a measure termed relative exploration, which is the amount of movement of the body divided by the minimum amount of movement required by perfect tracking (Methods). Under this definition, “1x” relative exploration indicates that the tracking trajectory traveled the same distance as the target trajectory. In the presence of additional exploratory movement as seen in Figure 2D, the relative exploration will exceed “1x”. Across the full fish behavior data set, we found a significant increase in relative exploration (Figure 3A upper row, Kruskal-Wallis test, *p* < 0.001, *n* = 21). This trend is reproduced with EIH which shows good agreement with the measured behavior, with significantly increased relative exploration as the signal weakens (Figure 3A lower row, Kruskal-Wallis test, *p* < 0.001, *n* = 18).

**Figure 3.**
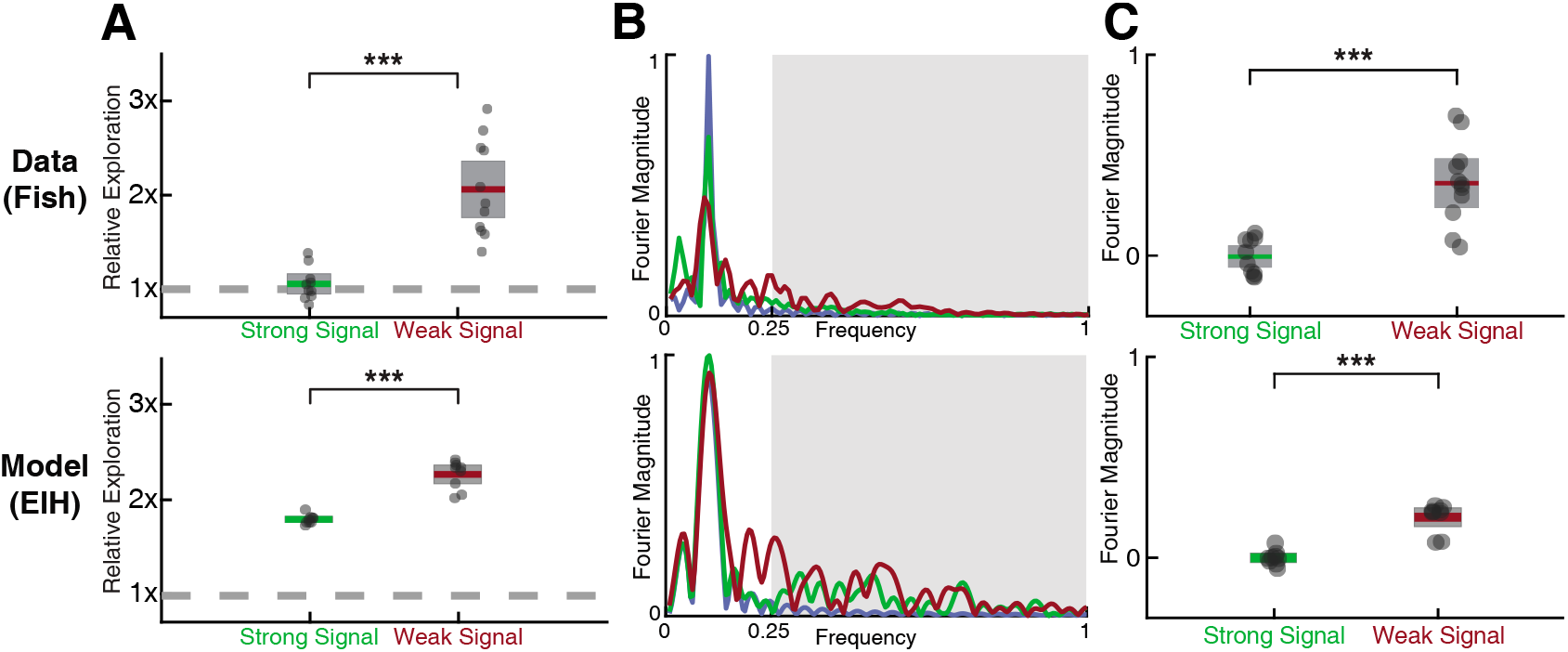
Fish behavior versus EIH predictions. (**A**) Relative exploration values (defined in text) for the fish and EIH trajectories under strong and weak signal conditions. Each dot represents a behavioral trial or simulation. EIH (bottom row) shows good agreement with behavioral data (upper row) as both have significantly higher relative exploration in the weak signal condition (Kruskal-Wallis test, *p* < 0.001, *n* = 21 for experimental data, and *p* < 0.001, *n* = 18 for EIH). (**B**) Representative Fourier spectra of the fish and EIH refuge tracking trajectories as seen in Figure 2C–D and F–I, with target trajectory (blue), strong signal condition (green), and weak signal condition (red). The frequencies above the frequency domain of the target’s movement are shaded; components in the non-shaded region are excluded in the subsequent analysis. The Fourier magnitude is normalized. (**C**) Distribution of the mean normalized Fourier magnitude within the sensing-related movement frequency band (gray shaded region) for strong and weak signal trials. These distributions are shown after subtraction by the sample mean of the strong signal data to emphasize the difference between strong and weak signal conditions. Each dot represents a behavioral trial or simulation. Significantly higher magnitude is found within the sensing-related movement band under weak signal conditions (Kruskal-Wallis test, *p* < 0.001, *n* = 21 for measured behavior, and *p* < 0.001, *n* = 18 for EIH). Asterisks indicate the range of *p* values for the Kruskal-Wallis test (* for *p* < 0.05, ** for *p* < 0.01, and *** for *p* < 0.001).

### The increase in exploratory movement is from sensing-related high frequency movement

To further characterize the sensing-related movement patterns and verify whether the increase in exploration is mainly due to these movements, we performed a spectral analysis of the fish’s tracking response. In Figure 3B we show the frequency spectrum of the refuge tracking trajectory shown in Figure 2C–D and F–I. Two frequency bands can be identified: 1) a baseline tracking band that overlaps with the frequency at which the target (the refuge) was moved; and 2) a sensing-related movement frequency band that accounts for most of the increased exploratory movements as signal weakens. The Fourier magnitude is significantly higher for the sensing-related movement frequency band under weak signal conditions when compared to strong signal conditions, for both the measured fish behavior (Figure 3C upper row, Kruskal-Wallis test, *p* < 0.001, *n* = 21) and EIH (Figure 3C lower row, Kruskal-Wallis test, *p* < 0.001, *n* = 18). This confirms that the significantly increased relative exploration reported in Figure 3A is primarily from sensing-related movements rather than baseline tracking movements.

### Sensing-related movements increase fish refuge tracking performance

A crucial issue to address is whether the additional sensing-related motions measured in the weak signal condition and predicted by EIH cause improved tracking performance. To answer this question, we constructed a filter to selectively attenuate only the higher frequency motion components without affecting the baseline tracking motion (Methods). Simulated weakly electric fish tracking trajectories in the weak signal condition—similar to that shown in Figure 2H—were filtered at increasing levels of attenuation. This led to a decrease in sensing-related body oscillations without affecting the baseline tracking motion (pre- and post-filtered trajectories: Figure 4—figure supplement 1). Filtered trajectories were then provided as the input to a sinusoidal tracking simulation in which the sensor moved according to the filtered trajectory. With respect to the full EIH sequence shown in Figure 2E, the final Trajectory Optimization step was removed and the trajectory was instead set to the filtered trajectory. The other elements of the EIH algorithm were held constant (Algorithm 1).

We show the results in Figure 4A in terms of relative tracking error, where 50% error means a departure from perfect tracking that is one-half the amplitude of the refuge’s fore-aft sinusoidal motion. Relative tracking error increases in proportion to the amount of sensing-related motion attenuation, from ≈50% with no attenuation to ≈75% with the highest attenuation we used. We then evaluated the distance from ergodicity, a dimensionless quantity that measures how well a given trajectory matches the corresponding EID distribution (Methods), for all the trajectories. We found that an increase in attenuation also leads to monotonically increasing distance from ergodicity. This indicates that the filtered trajectories are progressively worse at proportionally betting on information (Figure 4B). Figure 4C combines these two analyses, demonstrating that the distance from ergodicity is proportionate to tracking error.

**Figure 4.**
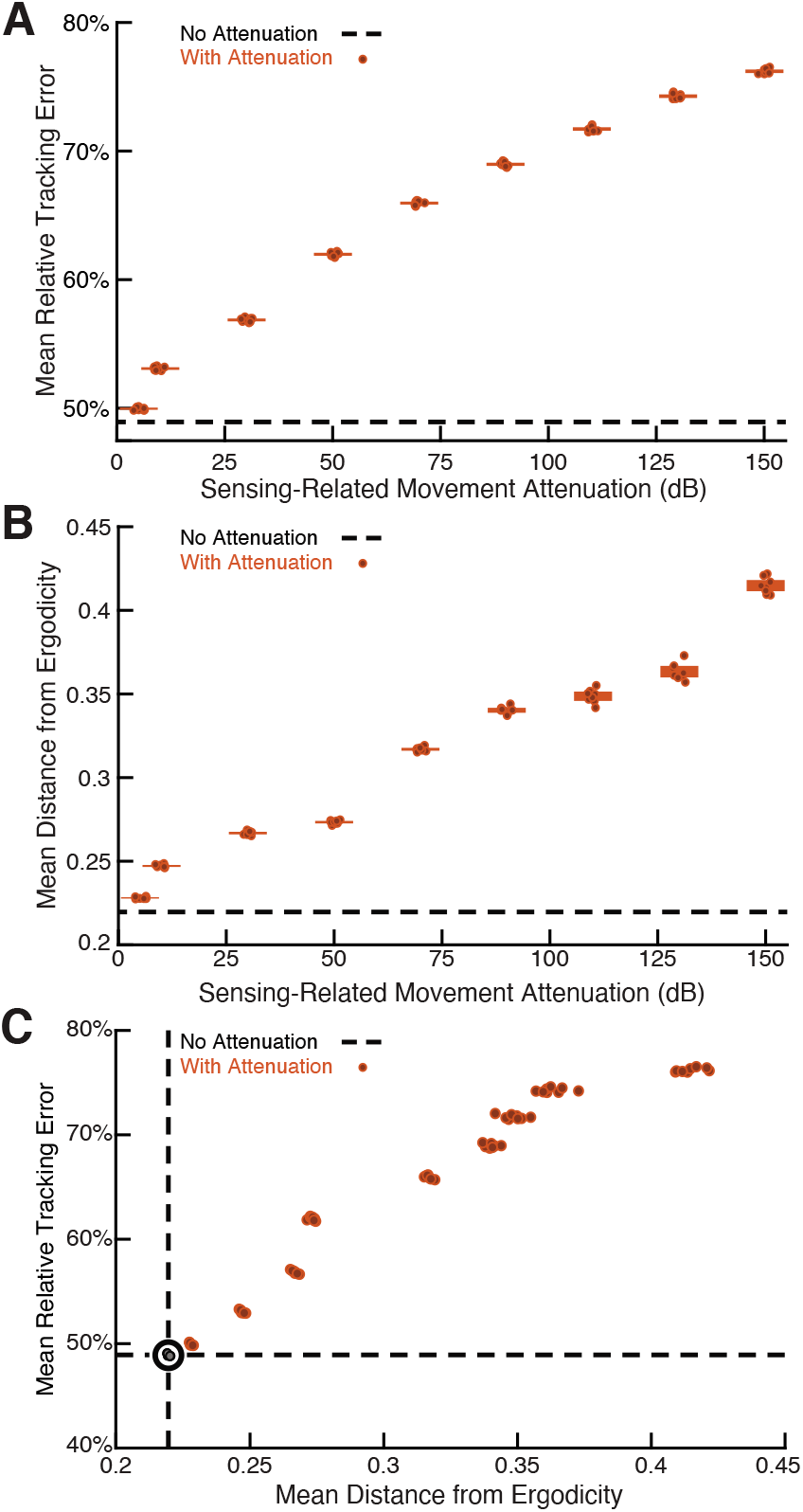
Sensing-related movements reduce tracking error. The full-body oscillation in the simulated EIH weak signal sensor trajectory similar to Figure 2H & I was gradually removed through a ramp increase of attenuation level (Methods) with 8 trials per condition (total *n* = 80) to establish the confidence interval. (**A**) Relative tracking error (% of amplitude of the target’s fore-aft movement, Methods) as sensing-related movement is attenuated. The line near 45% error shows relative tracking error for the original unfiltered trajectory (0 dB attenuation). The thickness of the horizontal bars represents the 95% confidence interval across the eight trials (individual dots) for each attenuation condition. For each attenuation level, the individual trial dots are plotted with a small horizontal offset to enhance clarity. The baseline tracking error with no attenuation is marked by the dashed black line. (**B**) Distance from ergodicity (Methods) as a function of sensing-related movement attenuation. Zero distance from ergodicity indicates the optimal trajectory that perfectly matches the statistics of the EID. As the distance increases from zero, the corresponding trajectory samples the EID further from the perfect proportional-betting ideal. The baseline data with no attenuation is marked by the dashed black line. (**C**) Relative tracking error plotted against distance from ergodicity across attenuation level. There is a clear positive correlation between tracking error and distance from ergodicity as sensing-related movements are diminished. The following figure supplements are available for figure 4: **Figure 4—figure supplement 1.** How sensing-related movements were attenuated for analyzing the impact of their diminishment.

### Error versus energy expenditure during fish refuge tracking

We estimated the mechanical energy needed to move the fish body along the measured trajectories in comparison to moving the body along the exact trajectory of the refuge and define the ratio between them as the relative energy. Under this definition, any motion beyond baseline tracking will lead to a higher than 1x relative energy. Electric fish are estimated to have needed significantly more mechanical energy during tracking in the weak signal condition compared to the strong signal condition (≈ 4x more, Figure 5A, Kruskal-Wallis test, *p* < 0.001, *n* = 21). We also examined in simulation howtrackingerror relates to the estimated mechanical energy expended on moving the body, starting with the unfiltered (EIH) trajectories and progressing through higher attenuation levels that gradually eliminate sensing-related movements. This was done by computing the relative energy for the simulation data shown in Figure 4A. We found that the tracking error decreased as the relative energy increased, with diminishing returns as the relative energy level neared that needed for the original unfiltered EIH trajectory (≈ 30 times the energy needed to move the body along the refuge trajectory, Figure 5B).

**Figure 5.**
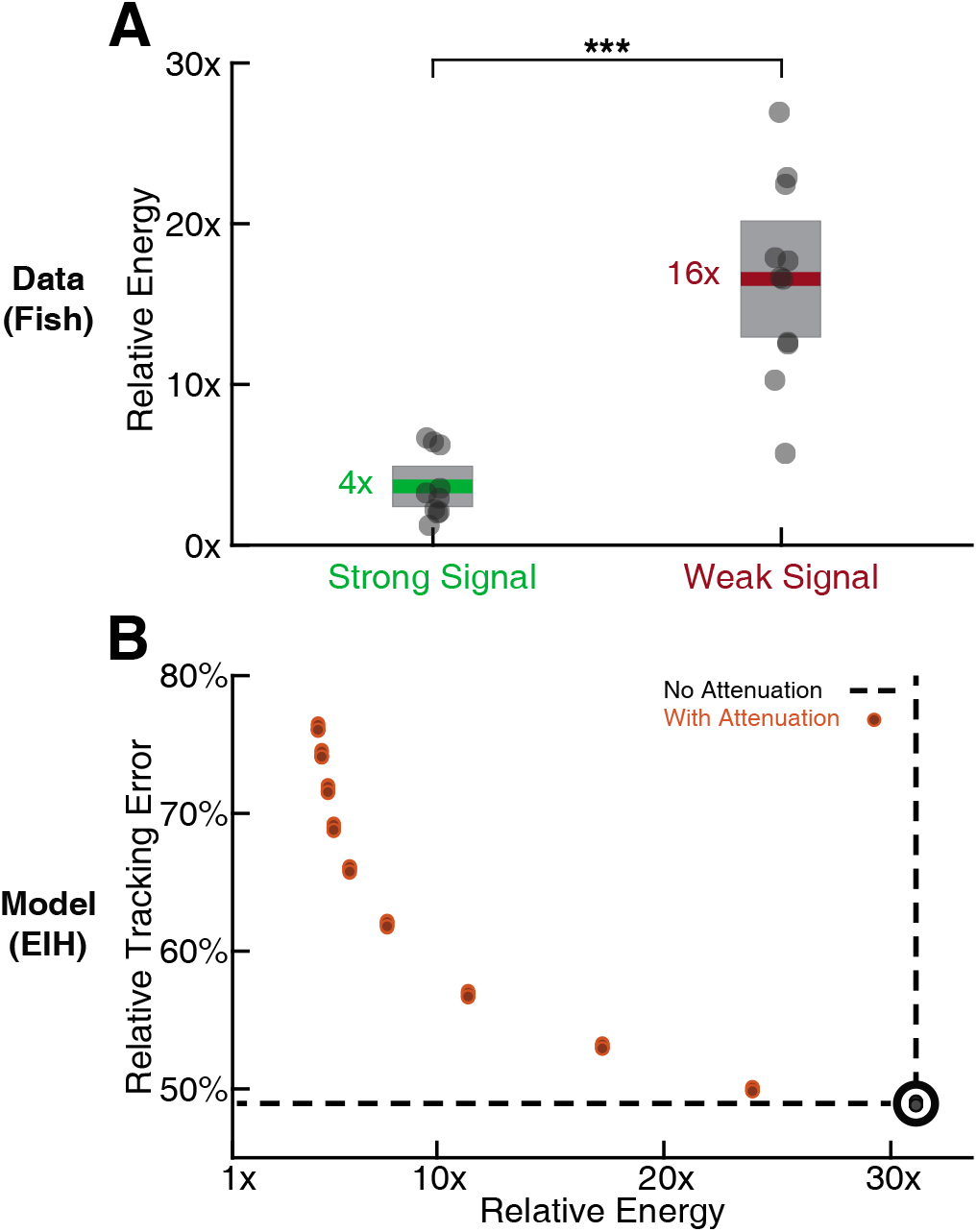
Relative energy for electric fish tracking behavior and EIH-generated behavior with attenuated body oscillations. (**A**) Relative energy (definition: text) used by the electric fish during refuge tracking behavior under strong and weak signal conditions. Trials are similar to those shown in Figure 2C–D. Weak signal conditions show a significantly higher relative energy as a result of the additional sensing-related movements (Kruskal-Wallis test, *p* < 0.001, *n* = 21). (**B**) Relative energy and relative tracking error for EIH simulations as sensing-related movements are progressively attenuated (data from Figure 4A-C, weak signal condition). As the sensing-related movements are attenuated, simulating investment of less mechanical effort for tracking, the relative energy decreases from 30x to less than 5x, but tracking error increases from 50% to around 75%. This plot suggests diminishing returns in tracking error reduction with additional energy expenditure beyond 30x. The lower bound near 4x is similar to the relative energy for the strong signal condition, and arises due to small disparities from the sinusoid that the refuge is following (the 1x path). Asterisks indicate the range of *p* values for the Kruskal-Wallis test.

### EIH predicts measured behavior across other species and sensory modalities

To evaluate whether EIH predictions can be generalized to the behavior of other animal species using other sensory modalities, we extended our analysis by running the EIH algorithm using previously published tracking data from other species. To allow comparison to the relative exploration analysis done for the fish, we selected datasets with strong and weak stimulus conditions (see Methods). In Figure 6—figure supplement 2 we also show an analysis of odor tracking in rats (***Khan et al., 2012***) but excluded from our analysis due to an insufficient number of trials. Figure 6 shows each species we considered other than hawkmoth flower tracking (analyzed separately). We include the electric fish for comparison. Below representative tracking trajectories for each species we show the corresponding EIH-predicted trajectory.

**Figure 6.**
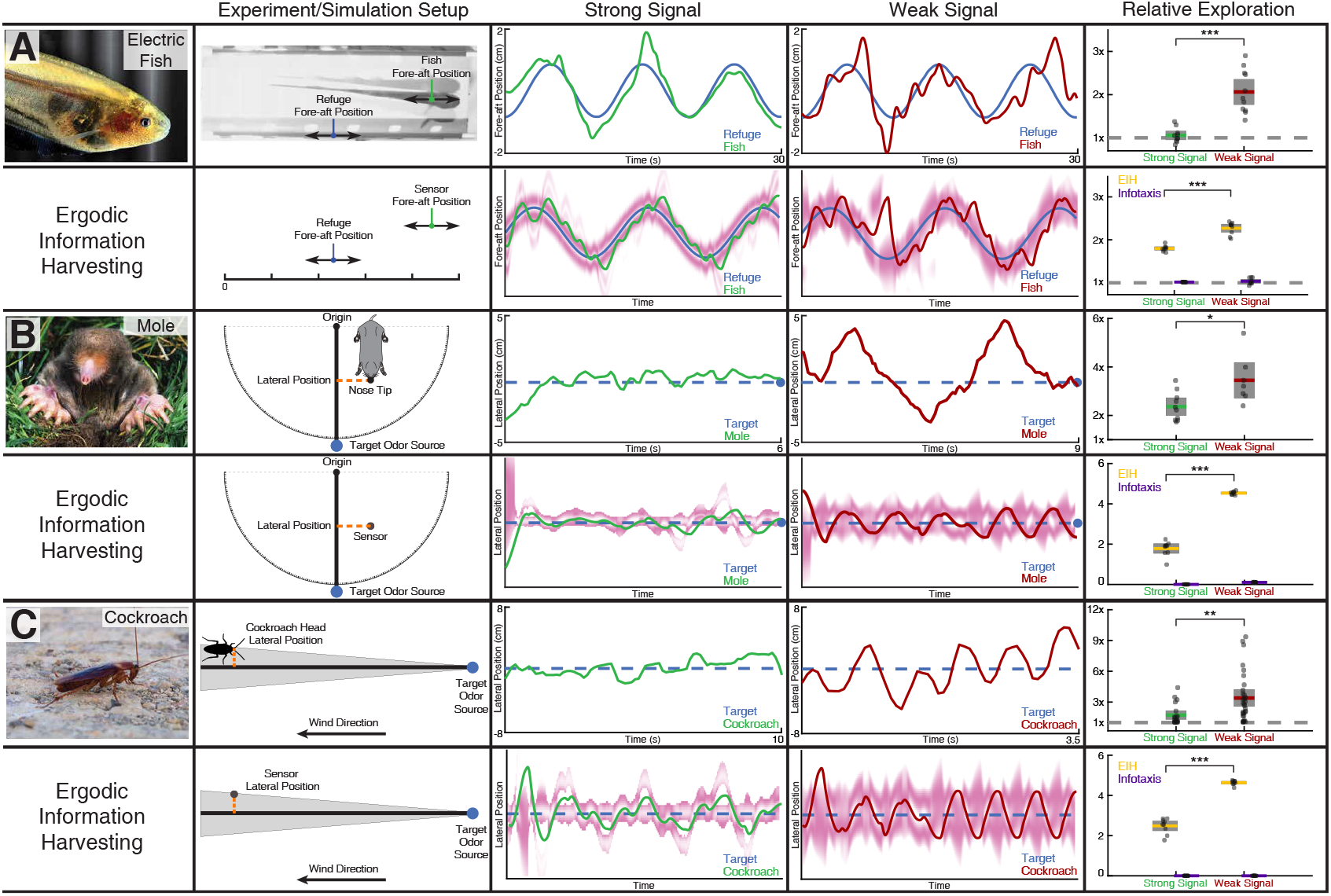
Trajectory comparison of animals tracking a target compared to EIH, and relative exploration across all trials. Three representative live animal trajectories above trajectories generated by the EIH algorithm, with their duration cropped for visual clarity. The moth data is not shown here due to the complexity of the prescribed target motion, but is shown in a subsequent figure. All EIH simulations were conducted with the same target path as present in the live animal data, using a signal level corresponding to the weak or strong signal categories (Methods). The EID is in magenta. Relative exploration across weak and strong signal trials (dots) is shown in the right-most column (solid line: mean, fill is 95% confidence intervals). Included for comparison is the relative exploration predicted by infotaxis (***Vergassola et al., 2007***). (**A**) The previously shown electric fish data to aid comparison to other species. As discussed, EIH agrees well with measured tracking behavior. Infotaxis (purple), in contrast, leads to hugging the edge of the EID, resulting in smooth pursuit behavior as indicated by the near 1x relative exploration. (**B**) The experimental setup and data for the mole was extracted from a prior study (***Catania, 2013***). During the mole’s approach to a stationary odor source, its lateral position with respect to the reference vector (from the origin to the target) was measured. Relative exploration was significantly higher under the weak signal conditions (Kruskal-Wallis test, *p* < 0.012, *n* = 17). In the second row, raw exploration data (lateral distance traveled in normalized simulation workspace units) are shown to allow comparison to simulation, as these are done in 1-D (Methods). EIH shows good agreement with significantly increased exploration for weak signal (Kruskal-Wallis test, *p* < 0.001, *n* = 18), while infotaxis leads to cessation of movement. (**C**)The experimental setup and data for the cockroach was extracted from a prior study (***Lockey and Willis, 2015***). The cockroach head’s lateral position was tracked and total travel distance was measured during the odor source localization task. Relative exploration is significantly higher for the weak signal condition (Kruskal-Wallis test, *p* < 0.002, *n* = 51). In the second row, we show that EIH raw exploration (as defined above) agrees well with measurements as the amount of exploration increased significantly under weak signal conditions (Kruskal-Wallis test, *p* < 0.001, *n* = 18), while infotaxis leads to cessation of movement. Asterisks indicate the range of *p* values for the Kruskal-Wallis test (* for *p* < 0.05, ** for *p* < 0.01, and *** for *p* < 0.001). Photo credit: Weakly electric fish (Original photo by Will Kirk, courtesy of Wikipedia, Creative Commons Attribution: Creative Commons, Attribution 2.5 Generic license); Mole (Original photo by Kenneth Catania, Vanderbilt University, courtesy of Wikipedia, Creative Commons Attribution-Share Alike 3.0 Unported license); Cockroach (Original photo by Sputniktilt, courtesy of Wikipedia, Creative Commons Attribution-Share Alike 3.0 Unported license). The following figure supplements are available for figure 6: **Figure 6—figure supplement 1.** Evolution of belief over time for the trials shown in Figure 6. **Figure 6—figure supplement 2.** Rat odor tracking behavior and EIH simulation. **Figure 6—figure supplement 3**. Measurements compared to prediction of EIH in two conditions where an animal needs to find the signal during tracking behavior. **Figure 6—figure supplement 4**. Systematic comparison between EIH and infotaxis in tracking a sinusoidally moving target. **Figure 6—figure supplement 5**. Sensitivity analysis on the ratio between control cost and ergodic cost in the objective function of trajectory optimization.

Figure 6B shows a representative trial of a mole engaging in a stationary odor source localization task (Methods). The behavioral data shows that the mole executes trajectories with significantly larger lateral oscillations under weak signal conditions (normal olfaction degraded by nostril blocking or crossing bilateral airflow, ***Catania 2013***) as summarized in the relative exploration plot (Kruskal-Wallis test, *p* < 0.001, *n* = 18). Figure 6C shows a trial of a cockroach localizing an odor source (Methods). Trials under weak signal conditions (normal olfaction degraded by trimming the olfactory antennae length, ***Lockey and Willis 2015***) show an increased amplitude of excursions from the odor track, which leads to a significant increase in relative exploration (Kruskal-Wallis test, *p* < 0.002, *n* = 51).

With respect to relative exploration, EIH shows good agreement with the measured behavior across these species, with significantly increased relative exploration as the signal becomes weak (Kruskal-Wallis test, *p* < 0.001, *n* = 18 for each species). Similarly good agreement was found for the increase in Fourier magnitude for sensing-related movement frequencies under weak signal conditions compared to strong signal conditions. This is shown in Figure 7 for the mole data (Figure 7E-H, Kruskal-Wallis test, *p* < 0.009, *n* = 17 for measured mole response and *p* < 0.001, *n* = 18 for EIH) and for the cockroach data (Figure 7I-L, Kruskal-Wallis test, *p* < 0.003, *n* = 51 for measured cockroach response and *p* < 0.001, *n* = 18 for EIH).

**Figure 7.**
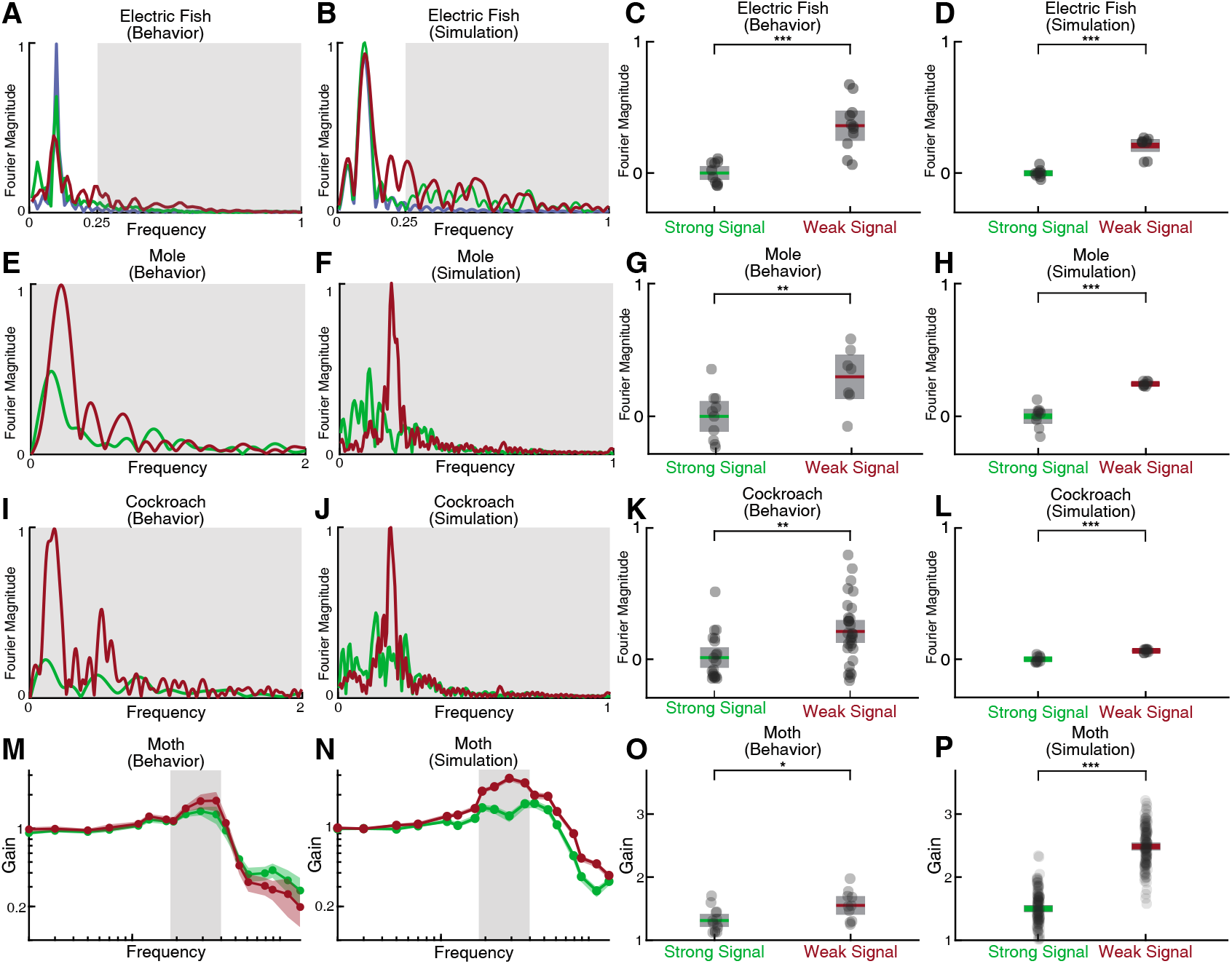
Spectral analysis of live animal behavior and simulated behavior. All the single trial Fourier spectra shown in A-B, E-F, and I-J are for the trials shown in Figure 6. (**A-B**) The already shown spectral analysis of the fish tracking data is included here for comparison, with target trajectory (blue), strong signal condition (green), and weak signal condition (red). The frequencies above the frequency domain of the target’s movement are shaded; components in the non-shaded region are excluded in the subsequent analysis. The Fourier magnitude is normalized. (**C-D**) The distribution of the mean normalized Fourier magnitude within the sensing-related movement frequency band (gray shaded region) for strong and weak signal trials, as shown before. (**E-F**) Representative Fourier spectra of the measured and modeled head movement of a mole while searching for an odor source (lateral motion only, transverse to the line between origin and target), plotted as in **A-B**. Because the target is stationary in contrast to the fish data, the entire frequency spectrum was analyzed. (**G-H**) Mean Fourier magnitude distribution, normalized to the strong signal condition to emphasize difference between the strong and weak cases. In the weak signal condition, there is significantly more lateral movement power under weak signal in both the animal data and simulation (Kruskal-Wallis test, *p* < 0.009, *n* =17 for behavior data, and *p* < 0.001, *n* = 18 for EIH simulations). (**I-J**) Fourier spectra of mole’s lateral trajectory, plotted the same as in **A-B**. (**K-L**) Mean Fourier magnitude distribution. Significantly higher power is found under weak signal in the behavior data and simulations (Kruskal-Wallis test, *p* < 0.003, *n* = 51 for behavior data, and *p* < 0.001, *n* = 18 for EIH simulations). (**M-N**) Bode magnitude plot (Methods) of a moth tracking a robotic flower that moves in a sum-of-sine trajectory (figure reprinted from ***Stockl et al. 2017***) and the corresponding simulation using the same target trajectory. Each dot in the Bode plot indicates a decomposed frequency sample from the first 18 prime harmonic frequency components of the flower’s sum-of-sine trajectory. A total of 23 trials (*n* = 13 for strong signal, and *n* = 10 for weak signal) were used to establish the 95% confidence interval shown in the colored region. Note the increase in gain in the midrange frequency region (shaded in gray) between the strong signal and weak signal conditions. The confidence interval for the simulation Bode plot is established through 240 trials (*n* = 120 for strong signal, and *n* = 120 for weak signal). The same midrange frequency regions (shaded in gray) are used for the subsequent analysis. (**O-P**) Mean midrange frequency tracking gain distribution for the moth behavior and simulation trials. Moth’s exhibit a significant increase in midrange tracking gain for the weaker signal condition (Kruskal-Wallis test, *p* < 0.02, *n* = 23), in good agreement with simulation (Kruskal-Wallis test, *p* < 0.001, *n* = 240). Asterisks indicate the range of *p* values for the Kruskal-Wallis test.

The last species we considered was hawkmoth tracking and feeding from a robotically controlled artificial flower (***Stockl et al., 2017***). In this case, the investigators used a complex sum-of-sines motion pattern for the artificial flower that is challenging to visualize in the same manner as we have plotted for the targets of other species. Instead we performed a spectral analysis that is similar to the Fourier magnitude analysis. We compared tracking when the moth was under high illumination (strong signal condition) and low illumination (weak signal condition) (Figure 7M-P, ***Stockl et al. 2017***). We analyzed the first 18 prime frequency components (up to 13.7 Hz) of both the moth’s response (Figure 7M, data from ***Stockl et al. 2017***, *n* = 13 for strong signal and *n* = 10 for weak signal) and simulation (Figure 7N, *n* = 120 for strong signal and *n* = 120 for weak signal), which is the same range used in ***Stockl et al. 2017***. We show the spectrum in Figure 7M-N as a Bode gain plot rather than Fourier magnitude since the target spectrum covers a wide frequency band including sensing-related movements (Methods). Consistent with previously reported behavior (***Stockl et al., 2017***), we found significantly increased mean tracking gain in the moth’s response within the mid-range frequency region relative to the strong signal condition (Figure 7O, Kruskal-Wallis test, *p* < 0.02, *n* = 23). This pattern is predicted by EIH simulations with the same sum-of-sine target trajectory (Figure 7P, Kruskal-Wallis test, *p* < 0.001, *n* = 240).

## Discussion

The body’s information processing and mechanical systems have coevolved to afford behaviors that enhance evolutionary fitness. Our theoretical approaches to these domains have proceeded along more independent tracks. Shortly after Shannon published his work on the information capacity of communication channels (***Shannon and Weaver, 1949***), his ideas were applied to visual perception (***Attneave, 1954**; **Barlow, 1959***) to describe efficient coding in the visual periphery. Since then, continual progress has been made in applying information theory to illuminate a host of problems in the coding and energetics of sensory signals from receptors to central nervous system processing (***Atick, 1992**; **Laughlin et al., 1998**; **Niven and Laughlin, 2008**; **Sengupta et al., 2010***). A parallel literature has matured analyzing animal motion (***Waldron et al., 2008**; **Srinivasan and Ruina, 2006**; **Ramdya et al., 2017**; **Nyakatura et al., 2019**; **Aguilar et al., 2016**; **Collins et al., 2005**; **Lee et al., 2008**; **McInroe et al., 2016**; **Sefati et al., 2013***). More recently these two areas are coming together in a growing literature that connects the information gathered through movement to the analysis of movement (***Cowan and Fortune, 2007**; **Rucci and Victor, 2015**; **MacIver et al., 2010**; **Sprayberry and Daniel, 2007**; **Stamper et al., 2012**; **Biswas et al., 2018**; **Yovel et al., 2010**; **Stockl et al., 2017**; **Fujioka et al., 2016**; **Yovel et al., 2011**; **Ghose and Moss, 2003, 2006**; **Hofmann et al., 2014**; **Bar et al., 2015**; **Nelson and MacIver, 2006**; **Bush et al., 2016**; **Yang et al., 2016***), but a general theory to bridge the gap between the information gained through movement and the energetics of movement is missing. Energy-constrained proportional betting is one such theory that is sufficiently general to invite application to a host of information-related movements observed in living organisms, while its algorithmic instantiation via EIH is sufficiently well-specified to generate testable quantitative predictions.

In the insects-to-mammals assemblage of animal species analyzed above, we observe gambling on information through motion, where the magnitude of the gamble is indexed by the energy it requires. EIH’s approach of extremizing a combination of ergodicity and energy generates trajectories that bet on information, exchanging units of energy for the opportunity to obtain a measurement in a new high-value location. For both measured and EIH-generated trajectories, a key change that occurs as sensory signals weaken is an increase in the rate and amplitude of the excursions from the mean trajectory, which we have quantified as an increase in relative exploration. Although the cause of these excursions in animals is poorly understood, the theory of energy-constrained proportional betting, and the implementation of the EIH algorithm based on that theory, can provide testable hypotheses.

First, the increase in the size of exploratory excursions in weak signal conditions arises with proportional betting because the EID spreads out in these conditions due to high uncertainty (for example, wider magenta bands in Figure 2I compared to 2G). Because a proportional betting trajectory samples proportionate to the expected information, as the expected information diffuses the excursions needed for its sampling will correspondingly increase in size.

Second, in EIH the spectral power profile of these excursions is related to the length of time interval for which a trajectory is generated (variable *T* in Figure 2, see Algorithm 1). One can consider this analogous to how far ahead in time an animal can plan a trajectory before changes in sensory information make planning irrelevant. For example, when tiger beetles see their prey, they execute a trajectory to the prey that is completed regardless of any motion of the prey after initiation of the trajectory. After each segment of running, if they have not caught their prey, they reorient their body toward it and enact a new trajectory and thereby gradually close the gap (***Gilbert, 1997***). In the strikes of the mottled sculpin, rather than a piece-wise open-loop ballistic strike, the entire strike is ballistic (***Coombs and Conley, 1997***). In contrast, prey strikes in electric fish are ballistic on time scales smaller than 115 ms but adaptive over longer time scales (***MacIver et al., 2001**; **Snyder et al., 2007***). In EIH, over the course of an enacted trajectory segment, changes in the expected information density due to new sensory observations similarly have no effect; these will only be incorporated in the generation of the next trajectory segment.

In EIH, perfect ergodicity is approached through the trade-off between the ergodic metric and the energy of movement within the prescribed trajectory time horizon *T*. As *T* asymptotically approaches infinite duration, the system will approach perfect ergodicity as the ergodic measure approaches zero. Conversely, as *T* asymptotically approaches zero duration, EIH will select a single direction to move to improve information. Changing *T* between these bounds will affect the frequency components of the sensing-related excursions in the context of exploring a bounded domain while searching for or tracking an object. This is because these excursions originate from the planned trajectory responding to the evolution of the EID, which is sampled at a rate of once every *T* prior to the synthesis of the next trajectory segment and assumed to be static in between.

For intuition on this point, again consider a fish locating a refuge along a line. As the fish moves to visit a region of high expected information in one direction, the unvisited locations in the other direction start to accumulate uncertainty—the belief distribution will begin to diffuse in those areas at a rate proportional to the noise level. This increase in uncertainty and its relationship to the observation model leads to an increase in expected information in those unvisited locations. This can be seen in Figure 2I—the deflection of the trajectory to investigate the fictive distractor present in the EID at the prior trajectory segment results in an increase in the EID in the opposite direction. After the sensor finishes the current trajectory segment of duration *T*, it then moves in the opposite direction to explore the unvisited regions with high expected information (Figure 6—figure supplement 1 shows the evolution of belief with respect to the trajectory segment boundaries). Because of these dynamics, a shorter *T* causes the sensor to react more quickly in response to changes in the EID and hence to higher frequency components within sensing-related movements. This same pattern, in combination with EIH’s tendency toward sampling across the EID, helps explain why sensing-related movements are often “oscillatory” (***Stamper et al., 2012***) or “zigzagging” (***Willy and Low, 2005**; **Webb et al., 2004***). The initial *T* (see Table 1) used for the behavior simulations was chosen to fit the frequency of sensing-related oscillations observed in the weakly electric fish refuge tracking data. The same value was applied to mole and cockroach trials, and reduced by a factor of five for the moth data due to the higher frequency content of the prescribed robotic flower movement.

**Table 1.**
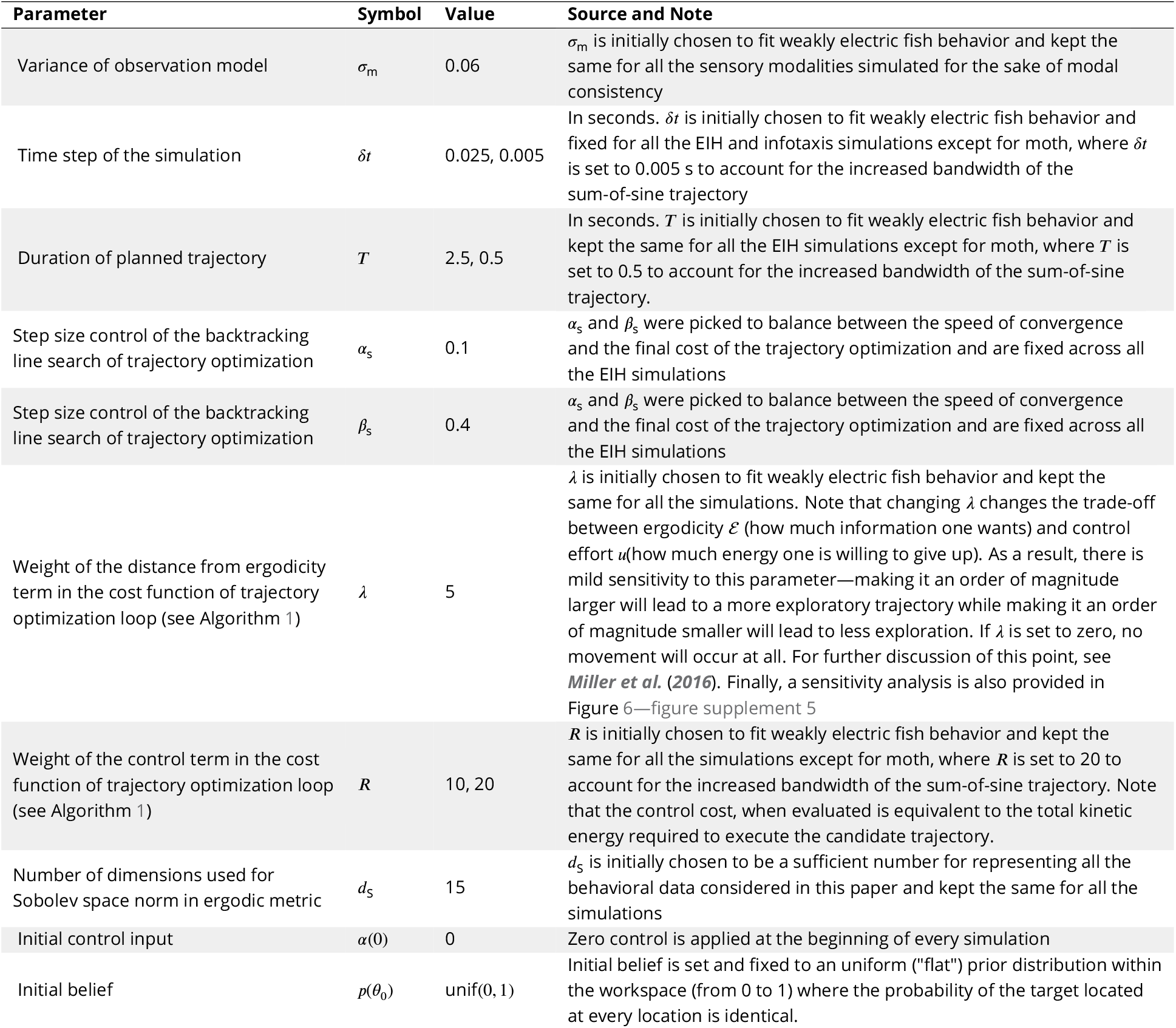
Parameters of EIH Simulation

| Parameter | Symbol | Value | Source and Note |
| --- | --- | --- | --- |
| Variance of observation model | $\sigma_m$ | 0.06 | $\sigma_m$ is initially chosen to fit weakly electric fish behavior and kept the same for all the sensory modalities simulated for the sake of modal consistency |
| Time step of the simulation | $\delta t$ | 0.025, 0.005 | In seconds. $\delta t$ is initially chosen to fit weakly electric fish behavior and fixed for all the EIH and infotaxis simulations except for moth, where $\delta t$ is set to 0.005 s to account for the increased bandwidth of the sum-of-sine trajectory |
| Duration of planned trajectory | $T$ | 2.5, 0.5 | In seconds. $T$ is initially chosen to fit weakly electric fish behavior and kept the same for all the EIH simulations except for moth, where $T$ is set to 0.5 to account for the increased bandwidth of the sum-of-sine trajectory. |
| Step size control of the backtracking line search of trajectory optimization | $\alpha_s$ | 0.1 | $\alpha_s$ and $\beta_s$ were picked to balance between the speed of convergence and the final cost of the trajectory optimization and are fixed across all the EIH simulations |
| Step size control of the backtracking line search of trajectory optimization | $\beta_s$ | 0.4 | $\alpha_s$ and $\beta_s$ were picked to balance between the speed of convergence and the final cost of the trajectory optimization and are fixed across all the EIH simulations |
| Weight of the distance from ergodicity term in the cost function of trajectory optimization loop (see Algorithm 1) | $\lambda$ | 5 | $\lambda$ is initially chosen to fit weakly electric fish behavior and kept the same for all the simulations. Note that changing $\lambda$ changes the trade-off between ergodicity $\mathcal{E}$ (how much information one wants) and control effort $u$ (how much energy one is willing to give up). As a result, there is mild sensitivity to this parameter—making it an order of magnitude larger will lead to a more exploratory trajectory while making it an order of magnitude smaller will lead to less exploration. If $\lambda$ is set to zero, no movement will occur at all. For further discussion of this point, see *Miller et al. (2016)*. Finally, a sensitivity analysis is also provided in Figure 6—figure supplement 5 |
| Weight of the control term in the cost function of trajectory optimization loop (see Algorithm 1) | $R$ | 10, 20 | $R$ is initially chosen to fit weakly electric fish behavior and kept the same for all the simulations except for moth, where $R$ is set to 20 to account for the increased bandwidth of the sum-of-sine trajectory. Note that the control cost, when evaluated is equivalent to the total kinetic energy required to execute the candidate trajectory. |
| Number of dimensions used for Sobolev space norm in ergodic metric | $d_s$ | 15 | $d_s$ is initially chosen to be a sufficient number for representing all the behavioral data considered in this paper and kept the same for all the simulations |
| Initial control input | $\alpha(0)$ | 0 | Zero control is applied at the beginning of every simulation |
| Initial belief | $p(\theta_0)$ | unif(0, 1) | Initial belief is set and fixed to an uniform ("flat") prior distribution within the workspace (from 0 to 1) where the probability of the target located at every location is identical. |

As gambling on information through motion involves a trade-off between increasing how well a trajectory approaches ideal sampling (ergodicity) and reducing energy expenditure, a useful quantity to examine is how tracking error changes with the energy expended on motion. To do so, we estimated the mechanical energy needed to move the body of the electric fish along the weak and strong signal trajectories, and found that weak signal trajectories required four times as much energy to move the body along as strong signal trajectories. In simulation, we examined how tracking error changes as more energy is invested in sensing-related movements. This analysis shows that the accuracy of tracking increases with the mechanical effort expended on sensing movements, with a 25% reduction of tracking error at the highest level of energy expenditure compared to the low energy case where sensing related movements are removed.

### Comparison to information maximization

Information maximization and EIH emphasize different factors in target tracking. First, if a scene is so noisy as to have fictive distractors or real distractors (physical objects similar to the desired target), this will generate more than one peak in the probability distribution representing the estimated target location. If the initial location at which information maximization begins is near the wrong peak, information maximization will result in going to that peak (such as to the distractor in Figure 1B) and stay at that location. With ergodic harvesting, information across a specified region of interest will be sampled in proportion to its expected magnitude (Figure 1C) constrained by the energy expenditure needed to do so. This leads to sensing-related movements that may, at first glance, seem poorly suited to the task: for example, if the distractor has higher information density, as it does in Figure 1C, then it will be sampled more often than the lower information density of the true target—but what is important here is that the true target is sampled at all, enabling the animal to avoid getting trapped in the local information maxima of the distractor. For information maximization, if 1) there is only one target of interest; 2) the EID is normally distributed; and 3) the signal is strong enough that false positives or other unmodeled uncertainties will not arise, then information maximization will reduce the variance of the estimated location of the single target being sought and direct movement toward the true target location. We interpret the poor agreement between infotactic trajectories and measured behavior as indicating that the conjunction of these three conditions rarely occurs in the behaviors we examined. Given that these behaviors were all highly constrained for experimental tractability, it seems likely to be even rarer in unconstrained three-dimensional animal behavior in nature.

The second area where these two approaches have different emphases is highlighted in cases where noise is dominating sensory input in high uncertainty scenarios as is common in naturalistic cases. Information maximization leads to a cessation of movement since no additional information is expected to be gained in moving from the current location (Figure 6—figure supplement 4A). Energy-constrained proportional betting will result in a trajectory which covers the space (Figure 6— figure supplement 4A): the expected information is flat, and a trajectory matching those statistics is one sweeping over the majority of the workspace at a density constrained by EIH’s balancing of ergodicity with energy expenditure. For information maximization, coverage can only be an accidental byproduct of motions driven by information maximization. Appropriate explorationexploitation tradeoffs emerge organically within EIH.

### Other interpretations of the behavioral findings

Fruit bats have been shown to oscillate their clicks from one side to the other of a target, rather than aiming their emissions directly at the target (***Yovel et al., 2010***). ***Yovel et al. (2010)*** suggested that this off-axis sensing behavior arises from the bat aiming a peak in the maximum slope of the signal profile (similar to the behavior of infotaxis, Figure 1 B). Given that small changes in direction of the sonar beam lead to large changes in echo at the location of maximum slope of the sonar beam distribution, ***Yovel et al. 2010*** then concludes that the bat is localizing the target optimally. This optimal localization hypothesis is supported by noting that the Fisher information with respect to the sonar beam angle of attack is approximately maximized at the locations where the bats direct the sonar beams. However, this analysis only provides an partial connection to information maximization, because the maximum slope of the sonar beam intensity distribution does not necessarily, or even typically, correspond to the a peak in the EID. The main difference between the two is that EID accounts for the belief distribution—that describes where the target might be and the uncertainty associated with that estimate—whereas the sonar beam distribution only coincides with the EID if the one assumes that the ground truth target is located at the peak of the sonar intensity. This means that the bats ‘know’ where the target is and are using sonar to maintain an estimate rather than globally searching for the target. This interpretation would indicate that the analyzed behavior in ***Yovel et al. 2010*** is in a late phase of the sonar emission behavior, where the goal is mainly about maintaining an already good estimate of target location. In contrast, EIH performs well in the early stages of search behavior, when the target location is very uncertain and the EID plays a dominant role in behavior. In all the EIH simulations presented here, for example, the simulation began with a uniform prior for the belief, the highest level of uncertainty about target location possible.

Another hypothesis is that active sensing movements arise from the animal adapting its closed-loop tracking gain response to a reduction in signal contrast (***Borst et al., 2005**; **Ghose and Moss, 2006**; **Maimon et al., 2010**; **Biswas et al., 2018***). However, this gain adaptation hypothesis is underspecified, in the sense that critical components are missing to formulate an algorithm that generates predictive trajectories conformingto the hypothesis. If gain adaptation is implemented with a Bayesian filter and a process is specified to generate oscillatory motion around targets according to the variance of the belief as a measure of uncertainty, then in the narrow context of a single target with no distractors (neither real nor fictive due to high uncertainty), such an algorithm can be tuned to behave similarly to EIH. However, in more realistic scenarios, there is no apparent mechanism to address real or fictive distractors, a capability of EIH we elaborate on further below. Further work is needed to test the differences between EIH and the gain adaptation hypothesis, or to determine whether gain adaptation is an implementation of EIH in specific, biologically relevant circumstances.

***Khan et al. (2012)*** show that in rat odor tracking behavior only about 12% of the trajectory qualifies as edge-tracking, suggesting that the rat’s zig-zagging trajectory is not centered on the edge of the trail—as predicted by the information maximization hypothesis—but rather on the middle of the odor trail, consistent with ergodic harvesting. They also introduced a model for odor tracking that instructs the sensor to move forward and lateral at a fixed velocity and make decisions to switch the direction of lateral movements based on specific events of sensor measurement and position. Although their model could in principle be adjusted to fit the trajectories of animal tracking under weak signal, their zig-zag sensing-related movements are explicitly programmed to appear based on ad-hoc strategies. This makes the model less generalizable and yields little insight into the underlying mechanism. In contrast, the sensing-related movements that emerge with EIH are not programmed but arise naturally in the manner described above. In addition, the Khan et al. model lacks the ability to address distractors, as shown in Figure 6—figure supplement 1 and Figure 2—figure supplement 2, since the movement strategy is not based on the belief or EID map, whereas EIH naturally provides coverage in these scenarios.

Finally, Rucci and Victor (***Rucci and Victor, 2015***) and Stamper et al. (***Stamper et al., 2012***) propose that active sensing movements are the outcome of an animal actively matching the spatial-temporal dynamics of upstream neural processing—a process by which the movement serves as a “whitening filter” (***Rucci and Victor, 2015***) or “high-pass filter” (***Stamper et al., 2012***). Sensing-related movements could be for preventing perceptual fading (***Kunapareddy and Cowan, 2018***), which has similarities to the high pass filter hypothesis in that motion is to counter sensory adaptation, a high pass filter-like phenomena. Although evidence for the perceptual fading hypothesis during tracking behaviors is lacking, EIH shows good agreement with animal behavior without any mechanism for sensory adaptation. Similar to the gain adaptation hypothesis, the high-pass filter hypothesis is also missing key components for trajectory prediction. Nonetheless, when implemented with the missing components, including a Bayesian filter and a feedback process that generates trajectories that match the desired spatial-temporal dynamics (***Biswas et al., 2018***), the high-pass and whitening-filter hypotheses do not conflict with EIH in single target cases with low uncertainty. This is because EIH also predicts a preferred frequency band for sensing-related movements that may match the preferred spectral power of upstream neural processing. However, in the context of multiple target scenarios, high uncertainty due to weak signal resulting in fictive distractors, or in the absence of any target, the same considerations apply to the high-pass and whitening filter hypotheses as were mentioned for the gain adaptation hypothesis. Further work is needed to test the differences between EIH and the high pass and whitening filter hypotheses, or to determine whether these are an implementation of EIH in specific, biologically relevant circumstances.

### Distractors and multiple targets

Given the above discussion, a capability of EIH that differentiates it from prior theories and that naturally arises from its distributed sampling approach is its ability to reject distractors and sample multiple targets. The live animal experimental data we analyzed did not feature either real distractors (here defined as objects having a distinguishably different observation model from that of the target) or multiple targets (multiple objects with identical observation models). Nonetheless, the EIH simulations suggest that what we are calling “sensing-related motions”—those movements that increase as sensory signals weaken—sometimes occur for rejection of fictive distractors while animals track single objects. A fictive distractor emerges when the current belief for the target’s location becomes multi-peaked; each peak away from the true target’s location is then a fictive distractor (illustrated by arrow in Figure 2I). Figure 6—figure supplement 1 shows the presence of these fictive distractors in the simulations of the fish, cockroach, and mole tracking behaviors, where we plot the belief rather than the EID. Fictive distractors arise in both the strong and weak signal conditions, but result in small amplitude excursions in the strong signal conditions because of the higher confidence of observations. In the simulated tracking behavior, the other source of sensing-related movements beside fictive distractor inspection is the increased spread of a (unimodal) EID as signals weaken, as earlier discussed.

False positive rejection has the signature of a digression from the nominal tracking trajectory; this digression ends when one or more samples have been received indicating there is no object present at the spurious belief peak, which then brings the believed target location back to some where closer to the true target position (Figure 6—figure supplement 1). In contrast, with a physical distractor, a digression should occur, but the incoming sensory signals support the hypothesis that the object being detected has a different observation model from that of the target, rather than the absence of an object. As none of our datasets include physical distractors, we investigated EIH’s behavior in this case with a simulated physical distractor. Figure 2—figure supplement 2 shows a simulated stationary physical distractor in addition to a stationary target. EIH is able to locate the desired target while rejecting the distractor. This result buttresses a finding in a prior robotics study, where we experimentally tested how EIH responded to the presence of a physical distractor and showed that an electrolocation robot initially sampled the distractor but eventually rejected it (Figure 8 of ***Miller et al. 2016***). In comparison, Figure 2—figure supplement 2 shows that infotaxis stalls as it gets trapped at one of the information maximizing peaks and fails to reject the distractor.

If, instead of a distractor and a target, EIH has two targets, the advantage of EIH’s sampling the workspace proportional to the information density is particularly well highlighted. A simulation of this condition is shown in Figure 2—figure supplement 1. EIH maintains good tracking with an oscillatory motion providing coverage for both of the targets. As seen in Figure 2—figure supplement 1, such coverage is not a feature of infotaxis, which gets stuck at the location of the first target and fails to detect the presence of the other target. A final case to consider is multiple targets with different (rather than identical) observation models. Tracking in such cases requires a simple adjustment to the calculation of the EID that we have explored elsewhere (Eq. 13 of ***Miller et al. 2016***).

While these preliminary simulations exploring how EIH performs with multiple targets and distractors are promising, it points to a clear need for animal tracking data in the presence of physical distractors or multiple targets (and in 2-D or 3-D behaviors: Appendix 6) in order to better understand whether EIH predicts sensory organ motion better than the gain adapatation or high-pass filtering theories in these cases.

### Biological implementation

The sensor or whole-body sensing-related movements we observe in our results is for proportional betting with regard to sensory system-specific EIDs—for electrosense, olfaction, and vision. To implement EIH, one needs to store at least a belief encoding knowledge about the target. The Bayesian filter update in EIH has the Markovian property, meaning that only the most recent belief is required to be stored. The EID, moreover, is derived from the belief and only used for every generated trajectory segment update, hence does not need to be stored. While the memory needs of EIH are low, trajectory synthesis requires computing the distance from ergodicity between candidate trajectories and the EID, a potentially complex calculation (Methods, Appendix 3). However, the complexity of our calculation may not be indicative of the complexity of implementation in biology. For instance, a recent study (***Stachenfeld et al., 2017***) suggests that a predictive map of future state is encoded in grid cells of the entorhinal cortex through spatial decomposition on a low-dimensionality basis set—a process similar to the calculation of the ergodic metric (Appendix 3). In weakly electric fish, electroreceptor afferents have power law adaptation in their firing rate in response to sensory input (***Drew and Abbott, 2006**; **Clarke et al., 2013***). This makes their response invariant to the speed of the target (***Clarke et al., 2013***) and hence similar to the simulated sensory input used to drive EIH. The power law adaptation also results in a very strong response at the reversal point during whole-body oscillations (Fig. 5 of ***Clarke et al. 2014***), a response generated by hindbrain-midbrain feedback loops (***Clarke and Maler, 2017***). Given the importance of an increased rate of reversals as signals become weaker in the fish tracking data and EIH, and EIH’s invariance to speed, the hindbrain area along with feedback loops to the midbrain are a worthwhile target for future work on the biological basis of EIH.

### Ergodic movement as an embodied component of information processing

As in the case of fixational eye movements (***Rucci and Victor, 2015***), a common interpretation of body or sensor organ movements away from the assumed singular goal trajectory is that this reflects noise in perceptual or motor processes. Ergodic harvesting presents a competing hypothesis: gathering information in uncertain complex environments means the system should be observed to move away from the singular goal trajectory. These excursions occur as predicted by EIH, including the possibility of multiple targets, and thus increase in size when uncertainty increases.

If an animal is at one peak of a multi-peaked belief distribution, what motivates it to move away from the current peak? The current peak already has sensor noise and other aspects of sensor physics incorporated, but misses other important sources of uncertainty. An occlusion may corrupt signal quality at a location otherwise predicted to have high target information. Other signal generators in the environment may emit confusing signals or the location may be contaminated by a fictive distractor arising from unmodeled uncertainty. Hence, the opportunity to visit another location in space that is statistically independent—yet contains a similar amount of predicted information—gives an animal an opportunity to mitigate unmodeled uncertainties through the expenditure of energy for movement. This is supported by experiments in human visual search suggesting that saccades are planned in a multi-stage manner for coverage of information towards the task-relevant goal rather than aiming for information maximization (***Yang et al., 2016; Hoppe and Rothkopf, 2019***). For example, a model to predict human visual scan paths found 70% of the measured fixation locations were efficient from an information maximization perspective, but there were many fixations (≈30%) that were not purely for maximizing information and attributed in part to perceptual or motor noise (***Yang et al., 2016***). We hypothesize that these apparently less efficient fixation locations are in fact the result of gambling on information through motion. It is also possible that motor noise may aid coverage in a computationally inexpensive manner.

The role of motion in this sensing setting is to mitigate the adverse impact of sensor properties. If, however, one is in an uncertain world full of surprises that cannot be anticipated, using energy to more fully measure the world’s properties makes sense. This is like hunting for a particular target in a world where the environment has suddenly turned into a funhouse hall of mirrors. Just as finding one’s way through a hall of mirrors involves many uses of the body as an information probe—ducking and weaving, and reaching out to touch surfaces—energy-constrained proportional betting predicts amplified energy expenditure in response to large structural uncertainties.

## Acknowledgments

We thank Mark Willis, Simon Sponberg, and Ken Catania for providing the original behavioral tracking data used for the studies we have cited. We thank the anonymous reviewers for many improvements as well as a suggestion on biological implementation. We thank Madhav Mani and Brennan Sprinkle for helpful discussions and feedback on an earlier draft. Funded by National Science Foundation IIS-1427419.

## Competing Interest

The authors declare no competing financial interest.

## Materials and Methods

### Electric fish electrosensory tracking

Three adult glass knifefish (*Eigenmannia virescens,* Valenciennes 1836, 8-15 cm in body length) were obtained from commercial vendors and housed in aquaria at ≈28°C with a conductivity of ≈100 *μS* cm^-1^. All experimental procedures were approved by the Institutional Animal Care and Use Committee of Northwestern University.

An experimental setup was built (similar to a prior study ***Stamper et al. 2012***) in which a 1D robot-controlled platform attached to a refuge allows precise movement of the refuge under external computer control. The refuge was a customized rectangular section, made by removing the bottom surface of a 15 cm long by 4.5 cm high by 5 cm wide PVC section (3 mm thick) and making a series of 6 openings (0.6 cm in width and spaced 2.0 cm apart) on each side. These windows provide a conductive (water) alternating with non-conductive (PVC) grating to aid electrolocation. The bottom of the refuge was 0.5 cm away from the bottom of the tank to ensure that the fish stays within the refuge. A high-speed digital camera (FASTCAM 1024 PCI, Photron, San Diego, USA) with a Nikon 50 mm f/1.2 fixed focal length lens was used to capture video from below the tank viewing up at the underside of the fish (Figure 2B). Video was recorded at 60 frames s^-1^ with a resolution of 1024 × 256 pixels. The refuge was attached to a linear slide (GL20-S-40-1250Lm, THK Company LTD, Schaumburg, USA), with a 1.25 m ball screw stroke and a pitch of 40 mm per revolution. The slide is powered by an AC servomotor (SGM-02B312, Yaskawa Electric Corporation, Japan) and servomotor controller (SGD-02BS, Yaskawa Electric Corporation, Japan). The refuge trajectory was controlled by a remote MATLAB xPC target with a customized Simulink model (MathWorks, Natick, USA).

Before each experimental session, individual fish were placed into an experiment tank (with identical water conditions) equipped with the moving refuge, high-speed camera and closed-loop jamming system (see below), and allowed to 2-4 hours to acclimate. Trials were done in the dark with infrared LEDs (*λ* = 850 nm) used to provide illumination for the camera. Each trial was 80 seconds long with the jamming signal only applied after the first 10 seconds and removed for the final 10 seconds. A total of 21 trials (*n* = 10 for strong signal and *n* =11 for weak signal) were used for this analysis. During each trial, the servomotor drives the refuge in a predefined 0.1 Hz sinusoidal fore-aft motion with an amplitude of 17 mm.

Video of electric fish refuge tracking was processed by a custom machine vision system written in MATLAB to obtain the fish head centroid and location of the refuge at 60 Hz. The *x* (longitudinal) position of the centroid of head of the fish was filtered by a digital zero-phase low-pass IIR filter with a cut-off frequency of 2.1 Hz and then aligned with the refuge trajectory. For all the completed trials (*n* = 21) across a total of 3 electric fish, the trajectory of both the fish and the refuge trajectory was used for the frequency domain analysis analysis (Figure 3). We used the Fourier transform to analyze the fish’s tracking response in the frequency domain. Trials with no jamming are categorized as the strong signal condition (*n* = 10, average trial duration 59.6 seconds). Trials with jamming (jamming amplitude ≥ 10 mA, see below) are categorized as the weak signal condition (*n* = 11, average trial duration 54.5 seconds). The cumulative distance traveled by the fish and refuge during refuge tracking was computed and denoted by *D_f_* and *D_r_,* respectively. Relative exploration was then defined as *D*_f_/*D*_r_.

### Closed-loop jamming system

During refuge tracking, the fish’s electric organ discharge (EOD) signal was picked up by two bronze recording electrodes and amplified through an analog signal amplifier (A-M Systems Inc, Carlsborg, USA) with a linear gain of 1000 and a passband frequency from 100 Hz to 1000 Hz. A data acquisition unit (USB 6363, National Instruments, Austin TX, USA) provided digitized signals used in a custom MATLAB script (MathWorks, Natick MA, USA) to identify the principal frequency component of the EOD. A sinusoidal jamming signal was then generated through the same USB interface using the digital to analog voltage output channel. The jamming signal’s frequency was set to be a constant 5 Hz below the fish’s EOD frequency as previously found most effective (***Bastian, 1987; Ramcharitar et al., 2005***). The synthesized jamming signal was sent to a stimulus isolator (A-M Systems Inc, Carlsborg WA, USA), which also converted the voltage waveform into a current waveform sent to two carbon electrodes aligned perpendicular to the EOD recording electrodes to avoid interference. The efficacy of the jamming stimulus was verified by examining its affect on the EOD frequency shown in Figure 2—figure supplement 3.

### Mole olfactory localization

Tracking data of blind eastern American moles (*Scalopus aquaticus,* Linnaeus 1758) locating a stationary odor source were digitized from a prior study (***Catania, 2013***). Three experimental conditions were used in the original study: one in which there was normal airflow (categorized in the strong signal condition), one where one nares was blocked (weak signal condition), and one where the airflow was crossed to the nares using an external manifold (also weak signal condition). Relative exploration was defined as the ratio between the cumulative 2-D distance traveled by a mole and a straight line from its starting position to the odor location *D*_mole_/*D*_line_, to allow direct comparison between strong and weak signal conditions despite differences in the mole’s initial position and target location across trials (Figure 6C). For the corresponding EIH simulation trials, we define relative exploration as the raw cumulative lateral distance traveled by the sensor since the simulation is done in 1-D.

### Cockroach olfactory localization

American cockroach (*Periplaneta americana,* Linnaeus 1758) odor source localization behavior data was acquired from a prior study (***Lockey and Willis, 2015***) through the authors. The cockroach’s head position was tracked during an odor source localization task. The same behavioral experiments were conducted with the odor sensory organ, the antennae, bilaterally cut to a specified length. The control group with intact antennae (4 cm in length) was categorized as the strong signal condition, and the 1 cm and 2 cm antenna-clipped groups were categorized as the weak signal condition. Only successful trials (*n* = 51, 20 strong signal condition trials and 31 weak signal condition trials) were included in the analysis. Relative exploration forthe cockroach data shown in Figure 6C is computed as the ratio of the cockroach’s total walking distance and the reference path length (the shortest path length from position at the start to the target) *D*_cockroach_/*D*_reference_ reported from the study (***Lockey and Willis, 2015***).

### Hawkmoth flower tracking

Hummingbird hawkmoths (*Macroglossum stellatarum,* Linnaeus 1758) naturally track moving flowers in the wind as they insert and maintain their proboscis in the nectary to feed, primarily driven by vision and mechanoreception (***Sponberg et al., 2015; Stockl et al., 2017***). In a prior study (***Stockl et al., 2017***), the hawkmoth’s flower tracking behavior was measured under various levels of ambient illumination while it fed from a nectary in a robotically controlled flower. The robotic flower moved in one dimension (lateral to the moth) in a predefined sum-of-sine trajectory composed of 20 prime multiple harmonic frequencies from 0.2 to 20 Hz. The moth’s lateral position was tracked and the Bode gain of the raw tracking data was acquired from a prior study (***Stockl et al., 2017***) through the authors and used in our analysis. A segment of the moth’s raw tracking trajectory is shown in Figure 1—figure supplement 1. We classified trials under high illumination (3000 lx, *n* = 13) as the strong signal condition, and trials under low illumination (15 lx, *n* = 10) as the weak signal condition.

### Non-technical description of EIH

The animal tracking simulations consist of several components, including simulation of the animal moving along a previously synthesized trajectory segment, simulating sensory observations, updating the simulated animal’s belief regarding the target’s location, computing the EID, and synthesizing the next trajectory segment by balancing ergodicity with the c
ost of movement (Figure 2E). This algorithmic implementation of energy-constrained proportional betting is built upon a framework we introduced in prior work for robotic tracking of stationary targets using Fisher information (***Miller et al., 2016***). The original algorithm was modified to track moving targets using entropy reduction as the information measure for better comparison to infotaxis, which also used this approach (***Vergassola et al., 2007***) (the results are insensitive to the choice of information metric; near identical results were obtained with Fisher Information). Code to reproduce these simulations, the empirical data, and a Jupyter Notebook stepping through how the EID is calculated, is available (***Chen et al., 2020***). For pseudocode of EIH and simulation parameters, see Algorithm 1 and Tables 1–2.

For each species, we model only one sensory system, the sensory system whose input was degraded through some experimental manipulation during the study. We model the sensory system as a point-sensor moving in one dimension (electrosense for fish, olfaction for mole and cockroach, and vision for moth). The sensory system is assumed to provide an estimate of location, modeled by the observation model which relates raw sensor measurements to the location of the target. Each sensor measurement also includes additive noise modeled by a Gaussian probability distribution with a variance determined by the specified signal-to-noise ratio (SNR)(see next section for more details about how sensory acquisition is simulated). Assuming this observation model and an initially uninformative (uniform) probability distribution of where the target is believed to be (this distribution is called the belief), EIH proceeds as follows: 1) For the current belief, derive the corresponding EID by calculating the answer to the question “how much information (quantified by entropy reduction) can we obtain by taking a new observation at this location?” for all possible locations (see the “EID Calculation” block in Figure 2E); 2) run the trajectory optimization solver to generate a trajectory segment with duration *T* (Table 1) that optimally balances energy expenditure against ergodicity with respect to the EID (the focus of this paper and the part of this animal simulation that is specific to this study; see the “Trajectory Synthesis & Optimization” block in Figure 2E); 3) execute the generated trajectory, allowing the sensor to make observations along it (see left half of the “Recursive Bayesian Filter” block in Figure 2E); 4) use the incoming observations to update the belief using a recursive Bayesian filter (***Thrun et al., 2005***). This is the step that updates where the simulated animal believes the target is located by taking into account new evidence and existing prior knowledge (see right half of the “Recursive Bayesian Filter” block in Figure 2E); 5) Check whether the termination condition has been reached (either running for a specified time or until the variance of belief is below a certain threshold), and if not, return to step 1. Video 2 shows these steps graphically for control of an bio-inspired electrolocation robot localizing a stationary target.

### Simulating Sensory Acquisition for Animal Tracking Simulations

For all analyzed animals, the body or sensory organ being considered is modeled as a point mass in a 1-dimensional workspace. The workspace is normalized to 0 to 1 for all the simulations. Each sensory measurement *V* is drawn from a Gaussian function that models the signal coming in to the the sensory system plus a zero-mean Gaussian measurement noise *ε* to simulate the effect of sensory noise:

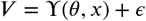

where *γ*(*θ,x*) denotes the Gaussian observation model function evaluated at position *x* given the target stimulus location *θ*

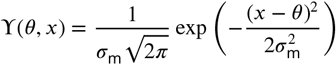

where *σ*_m_ is the variance of the observation model. Note that the observation model is not a statistical quantity, it is a deterministic map that relates the sensor reading (*θ* ∈ [0,1]) to the location of the target (*x* ∈ [0,1]). It should not be confused with the additive noise model which is a statistical quantity described by the Gaussian distribution. If we fix the target location *θ* at location 0.5 (center of the normalized workspace) and vary *x* to take continuous sensory measurements from 0 to 1, the resulting measurement versus location *x* will form a Gaussian (Figure 1D, upper inset).

The variance *σ*_n_ of *ε* is controlled by the signal-to-noise ratio (SNR) of the simulation:

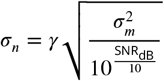

*σ*_n_ is the variance of the simulated sensory noise *ε*, and *γ* is a unity constant in units of normalized sensor signal unit per normalized workspace unit that converts *σ*_m_ (in normalized workspace units) on the RHS to normalized sensor signal units of the LHS term *σ*_n_. We used *σ*_m_ = 0.06 for all simulations (Table 1); The SNR values used for all the simulations is documented in Table 2. It should be noted that the values of SNR used in EIH simulations are only intended to relate qualitatively (“strong” or “weak” signal) to the actual (unknown) SNR of the animal’s sensory system during behavior experiments. To explore the impact of our assumptions regarding SNR, Figure 6—figure supplement 4 provides a sensitivity analysis showing how relative exploration varies as a function of SNR.

**Table 2.** Simulation Parameters Used For Each Figure

| Figure | Category | SNR (dB) | Initial Position | Target Trajectory | Biological Condition |
| --- | --- | --- | --- | --- | --- |
| Figure 1E & 2G, I & 3A-C | Weak Signal | $\leq 30$ | 0.4 | Sinusoid | N/A (simulation) |
| Figure 1F & 2F, H & 3A-C | Strong Signal | $\geq 50$ | 0.4 | Sinusoid | N/A (simulation) |
| Figure 2—figure supplement 1 | Weak Signal | $\leq 30$ | 0.4 | Stationary | N/A (simulation) |
| Figure 2—figure supplement 2 | Weak Signal | $\leq 30$ | 0.4 | Stationary | N/A (simulation) |
| Figure 4A-C & 5B | Weak Signal | $\leq 30$ | 0.4 | Sinusoid | N/A (simulation) |
| Figure 6A & 7B,D | Strong Signal | $\geq 50$ | 0.4 | Sinusoid | No jamming |
| Figure 6A & 7B,D | Weak Signal | $\leq 30$ | 0.4 | Sinusoid | $\geq 10$ mA jamming |
| Figure 6B & 7F,H | Strong Signal | $\geq 50$ | 0.2 | Stationary | Intact control |
| Figure 6B & 7F,H | Weak Signal | $\leq 30$ | 0.6 | Stationary | Single-side nares block & crossed airflow |
| Figure 6C & 7J,L | Strong Signal | $\geq 50$ | 0.475 | Stationary | 4 mm intact antenna |
| Figure 6C & 7J,L | Weak Signal | $\leq 30$ | 0.4 | Stationary | 1&2 mm bilaterally trimmed antenna |
| Figure 6—figure supplement 1 | Strong Signal | $\geq 50$ | 0.4 | Sinusoid | N/A (simulation) |
| Figure 6—figure supplement 1 | Weak Signal | $\leq 30$ | 0.4 | Sinusoid | N/A (simulation) |
| Figure 6—figure supplement 2 | Strong Signal | $\geq 50$ | 0.8 | Prescribed by study *Khan et al. (2012)* | Sham stitching |
| Figure 6—figure supplement 2 | Weak Signal | $\leq 30$ | 0.3 | Prescribed by study *Khan et al. (2012)* | Single-side nares stitching |
| Figure 6—figure supplement 3A | Weak Signal | $\leq 30$ | 0.9 | Prescribed by study *Khan et al. (2012)* | Single-side nares stitching |
| Figure 6—figure supplement 3B | Weak Signal | $\leq 30$ | 0.45 | Stationary | Single-side nares block |
| Figure 6—figure supplement 4 | Strong and Weak Signal | 10–55 | 0.4 | Sinusoid | N/A (simulation) |
| Figure 6—figure supplement 5 | Weak Signal | 20 | 0.4 | Sinusoid | N/A (simulation) |
| Figure 7N,P | Strong Signal | $\geq 50$ | 0.4 | Prescribed by study *Stockl et al. (2017)* | 3000 lux “high-light” |
| Figure 7N,P | Weak Signal | $\leq 30$ | 0.4 | Prescribed by study *Stockl et al. (2017)* | 15 lux “low-light” |

This generic Gaussian model of observation abstracts the process by which an animal relates afferent signals from sensory receptors to the estimated location of the target. The SNR of the observation model abstracts the effect of measurement uncertainty in the form of additive zero-mean Gaussian sensory noise. We used 10–30 dB as the weak signal condition and 50–70 dB as the strong signal condition due to the fact that relative exploration plateaus below 30 dB and beyond 50 dB SNR (Figure 6—figure supplement 4A). We only intend to use the relative change between high and low SNRs to simulate similar changes in the behavioral conditions of strong and weak signal trials.

The initial condition for all the simulation trials was a uniformly flat (uninformative) belief and an initial state of zero velocity and acceleration. To ensure uniformity, most of the simulation trials were set to have the exact same internal parameters except for SNR, which was changed across trials to compare trajectories in strong and weak signal conditions. The exception was the moth simulations, where we additionally used a smaller value for *T*, the duration of each planned trajectory segment, and a larger value of *R,* the weight of the control term in the objective function of the trajectory optimization (a larger *R* without changing the weight on the energy term *λ* means the trajectories are less energy constrained). A smaller *T* and larger *R* was needed due to the significantly higher bandwidth of the sum-of-sines trajectory prescribed for the robotic flower in the moth trials (Table 1).

### Quantifying information for the Expected Information Density (EID)

Consider the case of an animal tracking a live prey. Suppose that in open space the signal profile of the prey is similar to a 3-D Gaussian centered at its location. For a predator trying to localize the prey, sampling only at the peak of the Gaussian is problematic because while the signal is strongest at that location, it is also locally flat, so small variations in the prey’s location has little impact on sensory input. In contrast, any motion of the prey relative to the predator at the maximum slope of the signal profile will result in the largest possible change in the signal received by the predator, and therefore maximize the predator’s localization accuracy (Figure 1 D, the expected amount of information is maximal at the peak of the spatial derivative of the Gaussian profile).

Suppose at time *t* one has a probability distribution that is the belief *p*(*x*) about the value of *x*, for instance about the location of a particular chemical source, prey, or predator. In EIH, a Bayesian filter is used to optimally update *p*(*x*) from measurements, so *p*(*x*) evolves dynamically over time (***Kording and Wolpert, 2004***). The entropy of *p*(*x*), defined by ∑_*i*_*p*(*x_i_*)log(*p*(*x_i_*)) (where *i* is an index over a discretization of the domain), is the amount of information required to describe *p*(*x*) as a probability distribution. The entropy of a uniform or flat distribution is high—if it represents object location, it means an object could be at all possible locations in space and requires a lot of information to describe; while a narrow distribution for an object’s location can be described with very little information. The EID can be derived by simulating a set of possible sensing locations in the workspace, and for each location predicting the expected information gain by evaluating the reduction in entropy of the posterior with respect to the current prior (Appendix 4).

### Ergodicity

The ergodicity of a trajectory *s*(*t*) with respect to a distribution of the informativeness of sensing locations through space Φ(*x*) is the property that the spatial statistics of *s*(*t*)—the regions the trajectory visits and how often it visits them—matches the spatial distribution Φ(*x*). Technically, this is quantified by saying that a trajectory *s*(*t*) is ergodic with respect to Φ(*x*) if the amount of time the trajectory spends in a neighborhood 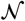 is proportional to the measure of that neighborhood 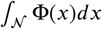 (Figure 1C). With a finite time horizon, perfect ergodicity is impossible unless one uses infinite velocity, which motivates a metric on ergodicity (***Scott, 2013***). A metric on ergodicity should be zero when a trajectory is perfectly ergodic and strictly positive and convex otherwise, providing a criterion that can be optimized to make a trajectory as ergodic as possible given the control cost constraint (see below). A standard metric used for comparing distributions is the Sobolev space norm, which can be computed by taking the spatial Fourier transform of Φ(*x*) and *s*(*t*) (see below). This metric is equivalent to other known metrics such as those based on wavelets (***Scott, 2013***). We can generate an ergodic information harvesting trajectory by optimizing the trajectory with respect to the ergodic metric (***Miller et al., 2016**),* often with real-time performance (***Mavrommati et al., 2018**),* both in deterministic and stochastic settings (***De La Torre et al., 2016***).

### Balancing energy expenditure and proportional betting

In EIH, candidate trajectories are generated (step 2 in the paragraph above) by minimizing the weighted sum of 1) the ergodic metric, which quantifies how well a given trajectory does proportionally betting on the EIH; and 2) the square norm of the control effort. Note that mass is implicitly included in the weighted sum. Optimizing the ergodic metric alone forms an ill-posed implementation as this implies that energy consumption is not bounded. This is equivalent to a situation where the energetic cost of movement is zero, with a consequent movement strategy of sensing everywhere. This is unlikely to be a reasonable movement policy for animals to maximize their chance of survival. Similarly, EIH is not minimizing energy either, as the energy minimizing solution alone is to not move at all. More realistically, when animals have a limited energy budget for movement, the control cost term should be added to impose a bound for energy consumption for a given trajectory. In the first-order approximation of the kinematics of motion of animals, the control cost is defined by the total kinetic energy required to execute the input trajectory (Algorithm 1). In our study, the control cost is not intended to explicitly model the energy consumption of any particular animal used in the study. It is used, however, to represent the fact that energy is a factor that animals need to trade-off with information while generating trajectories for sensory acquisition. The trade-off between ergodicity and the energy of motion is represented by *R/λ*, where *R* is the weight on the control cost and *λ* is the weight on the distance from ergodicity (Table 1). We used a value of *R/λ* = 2, resulting in a relative exploration value of around 2. The variation in relative exploration with an order of magnitude change in *R/λ* from 1 to 10 is 2.5 to 1.5 (Figure 6—figure supplement 5).

### Behavioral trajectory simulations

It is worth noting that in EIH, the animal’s tracking behavior is hypothesized to be the outcome of a dynamical system, the result of forces and masses interacting, rather than sample paths of a random process—the traditional venue for ergodicity and entropy to play a role in analysis. However, we discuss the possibility that sense organs are moved stochastically in Appendix 5. When used to simulate behavioral trajectories, EIH was reconfigured to use the prescribed stimulus path from the corresponding live animal experiment as the target trajectory (Figure 6). The simulated sensor’s initial position was set to match the animal’s starting location. To simulate the effect of a weak sensory signal, the SNR was reduced in the respective trials to simulate the effect of increased measurement uncertainty. Other than target trajectory, initial position, and SNR, the simulation parameters were the same across all simulations except for moth trials, where *T* (the duration of each planned trajectory segment) and *R* (the weight for the control cost term in the trajectory optimization objective function) are adjusted to better fit the increased bandwidth in the prescribed sum-of-sine target path (Table 2).

### Sensing-related movement attenuation simulations

The simulated electric fish tracking response under weak signal condition is filtered through zerophase IIR low-pass filters with different stop band attenuations (Figure 4—figure supplement 1A). These filters are configured to pass the low frequency tracking band within ≈0.2 Hz (target motion is a sinusoid in 0.1 Hz). This configuration allows effective removal of higher frequency sensing-related movements without affecting the baseline tracking response. The effect of the sensing-related motion filter is parameterized by the stop band attenuation at 1.5 Hz. The sensing-related movement magnitude can be systematically deteriorated by controlling the stop band attenuation while maintaining intact baseline tracking (Figure 4—figure supplement 1A-C).

The raw simulated weak signal tracking trajectory is first filtered by the sensing-related movement filter at stepped attenuation levels from 5 dB to 150 dB. The filtered trajectory is then prescribed to a tracking-only simulation where the sensor is instructed to move along the predefined input trajectory, take continuous sensor measurements, and use these to update the belief and EID. The distance from ergodicity is then evaluated based on the trajectory segment and simulated EID in the same way as for the other behavior simulations. Tracking performance is evaluated by comparing the sensor’s best estimate of the target’s position over time based on its belief and the ground truth.

### Infotaxis simulations

The original infotaxis algorithm (***Vergassola et al., 2007***) was adapted for 1-D tracking simulations. The infotaxis algorithm computes the EID in the same way as EIH, but differs in the movement policy once the EID is computed. The sensor considers three movement directions from its current position—left, right, or stay—at every planning update. The sensor follows the infotaxis strategy by choosing a movement direction that will maximize the EID and then takes samples along the chosen direction to update the Bayesian filter (***Thrun et al., 2005***) and consequently the EID for the next planning iteration. The parameters of the infotaxis simulation are kept the same as for EIH to allow direct comparison.

### Energetics

We analyzed how the additional movement for tracking in weak signal conditions affected energy use for electric fish (Figure 5). We estimated the net mechanical work required to move the fish along the observed tracking trajectory by first computing the instantaneous power *P*(*t*) = *F*(*t*)*v*(*t*) of tracking at every timestamp. The net force *F*(*t*) was estimated by applying Newton’s law *F* = *ma* with the estimated body mass *m* (from ***Postlethwaite etal. 2009***). Finally, the total mechanical work done by the fish is the integral of the instantaneous power over time *∫_t_P*(*t*)*dt*. The effect of added mass was included using equations previously developed (***Postlethwaite et al., 2009***).

The relative energy was defined as the total mechanical work of moving the fish along the tracking trajectory divided by the work of moving the fish along the trajectory of the target (the refuge). A relative energy of “1x” therefore indicates that moving the fish along the tracking trajectory required the same energy as moving it along the path that the moving refuge took.

### Spectral analysis

The frequency response of electric fish, mole, cockroach, and moth tracking and simulation data were analyzed using the Fourier transform. The magnitude frequency response data was used in Figure 3B-C and 7A-L. For the 2-D trajectories of mole and cockroach, the lateral tracking response was analyzed separately alongside the 1-D EIH lateral tracking simulation (Figure 7E-L). Because our simulations assume a normalized workspace dimension of 0 to 1, the spectral analyses are shown with normalized Fourier magnitudes and are only intended to provide a qualitative link between EIH and animal behavior, rather than matching the units of the original live animal trajectories. For the moth, since the sum-of-sine stimulus covers a wide frequency range that includes the frequencies of the sensing-related movements, the tracking response of moth behavior and simulation is shown in the form of a Bode gain plot (Figure 7M-N) instead of Fourier magnitude to visualize both the frequency spectrum of motion and relative exploration of each tracked frequency component. A Bode gain plot shows the magnitude of the frequency response of the tracking trajectory normalized by the stimulus for a wide range of frequencies. A gain of 1 for any particular frequency indicates the moth (or simulated sensor) responded with the same amplitude as the sum-of-sine stimulus at that frequency. The averaged Fourier magnitude and Bode gain were computed by taking the mean of the Fourier magnitudes or Bode gain within the sensing-related movement frequency window marked by the shaded area of the spectrum plots shown in the first columns of Figure 7. For electric fish, the sensing-related movement frequency window is identified as high frequency components outside of the baseline tracking response frequency range (Figure 3B-C and 7A-B). For the mole and cockroach, because the target is stationary and hence there is no baseline tracking frequency, the entire frequency spectrum of the tracking response was used for computing the statistics.

### Quantification and statistical analysis

The Kruskal-Wallis one-way ANOVA test was used for the statistical analysis of relative exploration (Figure 3 and 6) and spectral power of tracking (Figure 3 and 7). Each trial of weakly electric fish, mole, cockroach, and moth behavior as well as their corresponding simulations were considered independent. Kruskal-Wallis is non-parametric and hence can be applied to test for the significance of relative exploration even though it is a ratio distribution.

The Pearson correlation coefficient and the 95% confidence interval of its distribution were calculated in Figure 6—figure supplement 4B based on data from Figure 6—figure supplement 4A. The mean and 95% confidence interval was computed for Figure 3, 6, and 6—figure supplement 4.

### Data and software availability

All data and code needed to reproduce our results is available (***Chen et al., 2020***), as well as an interactive Jupyter notebook tutorial on computing the EID. Algorithm 1 provides psuedocode, and Tables 1–2 provide the corresponding simulation parameters for the EIH algorithm. Finally, Video 1 provides a video explainer of the theory of EIH, and Video 2 applies it to controlling an underwater electrolocation robot.

---

**Figure 1—figure supplement 1.**
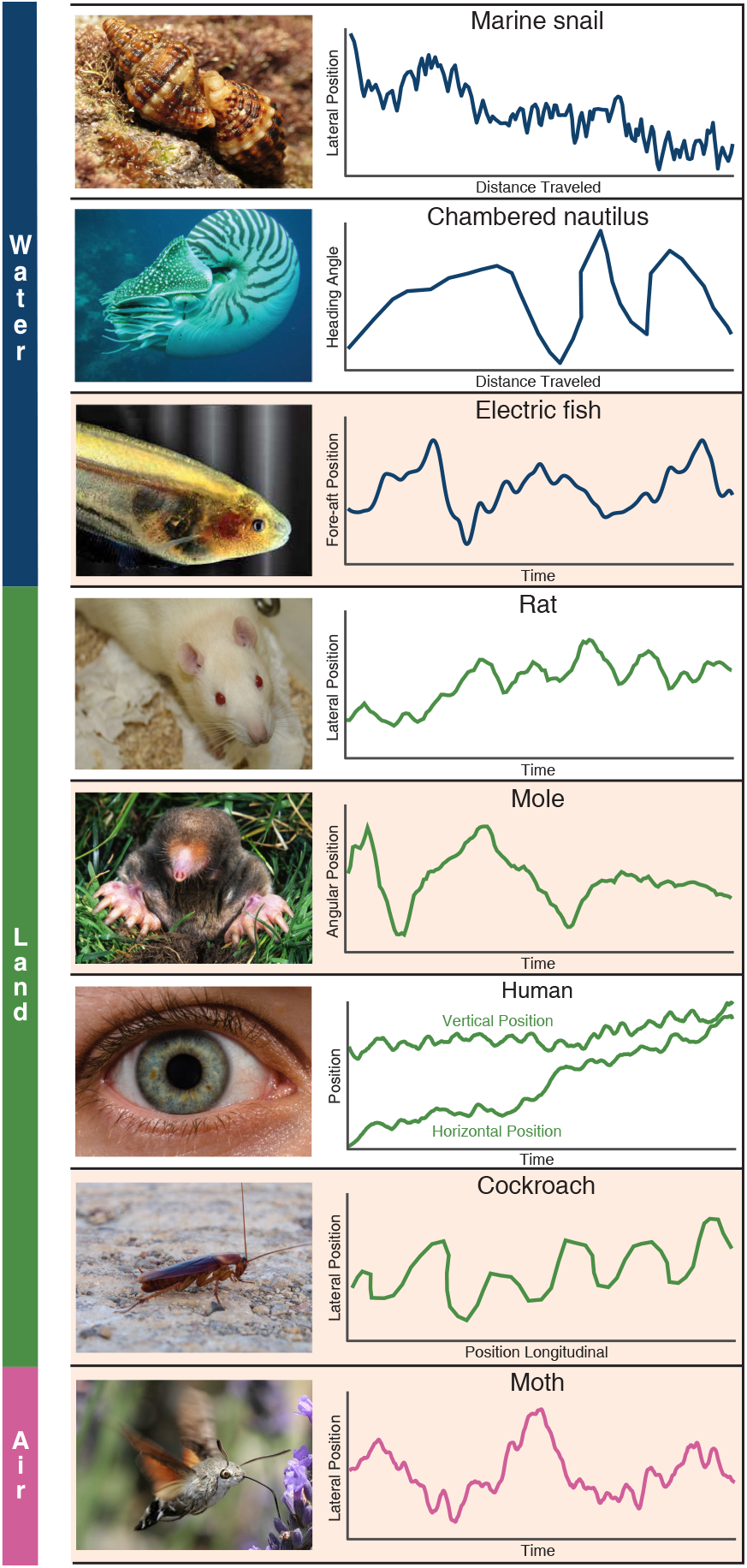
Whole-body or sensory organ small-amplitude motions are ubiquitous as animals track targets. Here we show siphon casting behavior in the marine snail (body, ***Ferner and Weissburg 2005***), cross-current swimming in the Chambered nautilus (body, ***Basil et al. 2000***), whole-body oscillations in electric fish (body position, ***Stamper et al. 2012***), zigzagging motions in the rat (nose, ***Khan et al. 2012***), back and forth searching in the mole (nose, ***Catania 2013***), fixational eye movements in humans (eye, ***Rucci and Victor 2015***), the zigzagging walk of the cockroach (head, ***Willis and Avondet 2005***), and flower tracking motions in the moth (head, ***Stockl et al. 2017***). These sensing-related movements occur with striking consistency across animals using different sensory modalities and within different physical environments. We propose these movements arise as a form of gambling on information through motion, and show evidence in support of this claim from the four species highlighted (electric fish, mole, cockroach, and moth). Photo credit: Marine snail (Original photo by Katja Schulz, courtesy of Wikipedia, Creative Commons Attribution: Creative Commons, Attribution 2.0 Generic license); Chambered nautilus (Original photo by Profbergert, courtesy of Wikipedia, Creative Commons Attribution-Share Alike 3.0 Unported license); Weakly electric fish (Original photo by Will Kirk, courtesy of Wikipedia, Creative Commons Attribution: Creative Commons, Attribution 2.5 Generic license); Rat (Original photo by Jean-Etienne Minh-Duy Poirrier, courtesy of Wikipedia, Creative Commons Attribution-Share Alike 2.0 Generic license); Mole (Original photo by Kenneth Catania, Vanderbilt University, courtesy of Wikipedia, Creative Commons Attribution-Share Alike 3.0 Unported license); Human (Original photo by ROTFLOLEB, courtesy of Wikipedia, Creative Commons Attribution-Share Alike 3.0 Unported license); Cockroach (Original photo by Sputniktilt, courtesy of Wikipedia, Creative Commons Attribution-Share Alike 3.0 Unported license); Moth (Original photo by IronChris, courtesy of Wikipedia, Creative Commons Attribution-Share Alike 3.0 Unported license).

**Figure 2—figure supplement 1.**
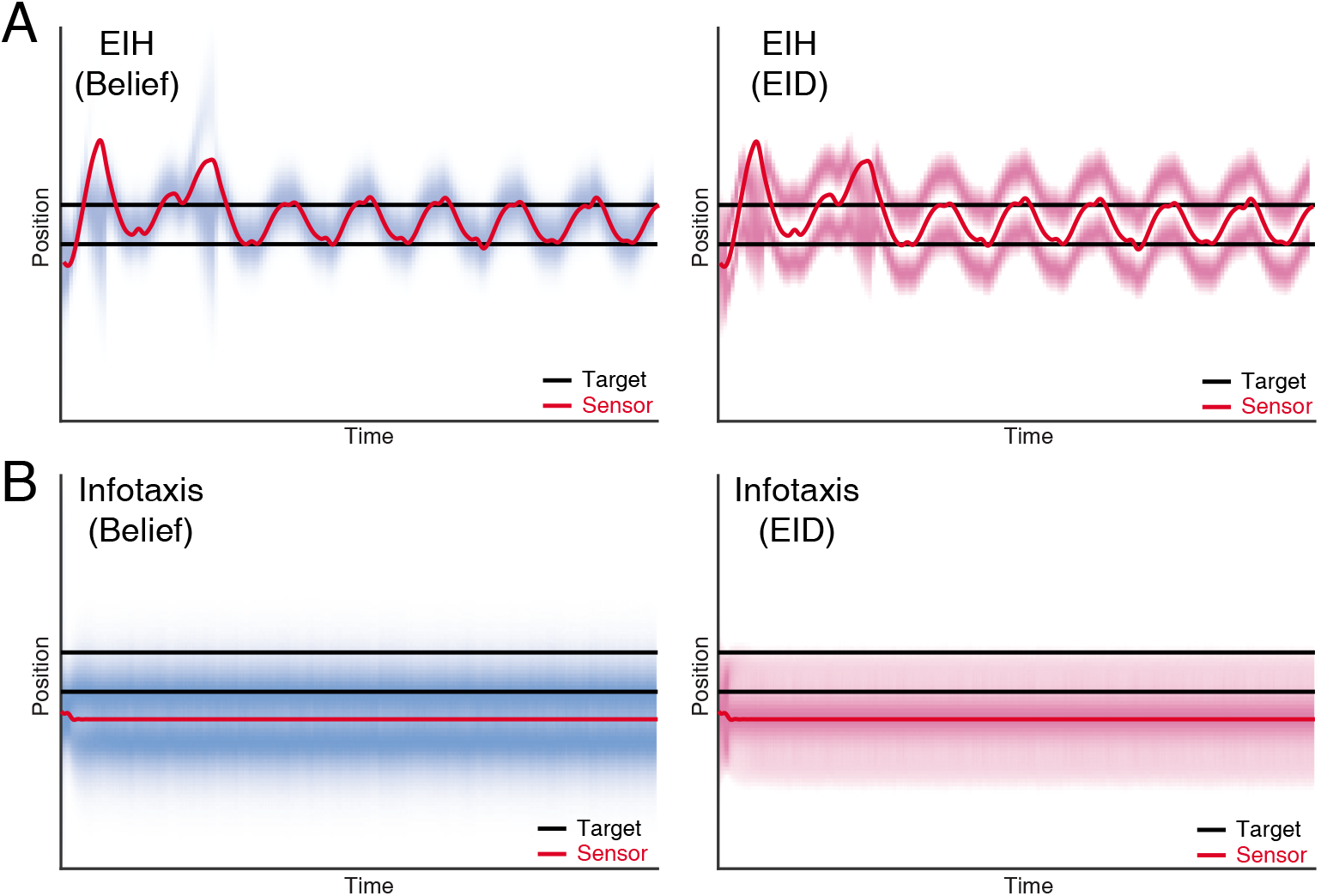
Dual target tracking simulation with EIH and infotaxis. In this simulation, two identical targets (with the same observation model) are present in the workspace, indicated by the two blue lines. To help visualize the outcome, the belief (left panel) and EID (right panel) distribution over time are rendered with blue and magenta color overlay, respectively. In both cases, we use identical SNR levels corresponding to a weak signal condition (30 dB, see Table 2). (**A**) EIH tracks both targets with an oscillatory motion. In addition, belief peaks corresponding to the top and bottom targets are clearly visible in the left panel. (**B**) Infotaxis gets trapped at one of the information maximizing peaks, as seen in the right panel. This leads to cessation of movement and failure to detect the other target on the top (left panel).

**Figure 2—figure supplement 2.**
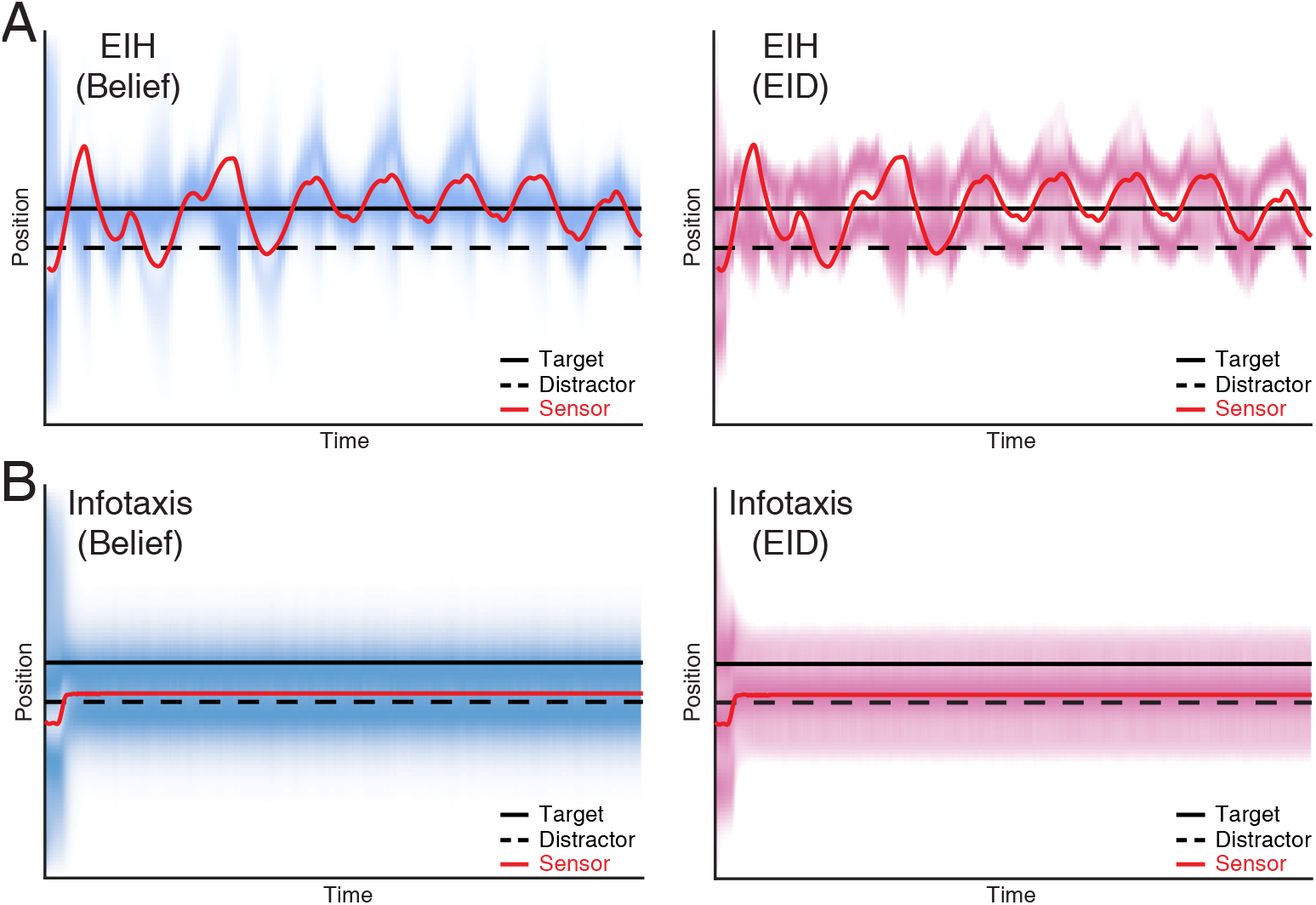
Single target tracking simulation with EIH and infotaxis in the presence of a simulated physical distractor. In this simulation, a single target and a physical distractor coexist in the workspace. The simulated physical distractor has a different observation model that leads to a different measurement profile when compared to the desired target. To help visualize the outcome, the belief (left panel) and EID (right panel) distribution over time are rendered with blue and magenta color overlay, respectively. In both cases, we use identical SNR levels corresponding to a weak signal condition (30 dB, see Table 2). (**A**) EIH is able to find the desired target (the continuous peak along the desired target location in the left panel) and rejects the distractor (no continuous peak along the distractor location in the left panel). (**B**) Infotaxis gets trapped at one of the information maximizing peaks, as seen in the right panel. Although it does find the desired target, it fails to reject the distractor, as seen in the continuous peak near the distractor in the left panel.

**Figure 2—figure supplement 3.**
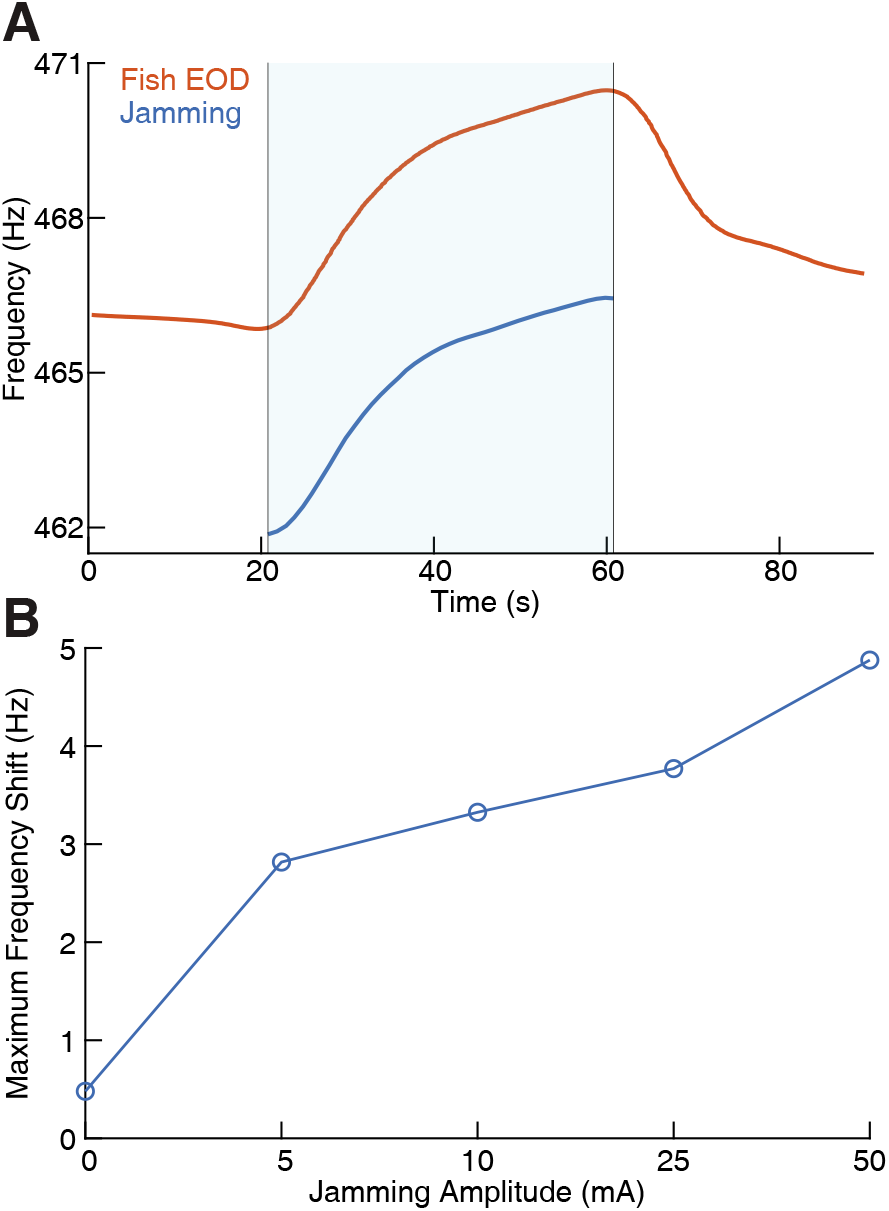
Effect of jamming and how it varies with jamming intensity. (**A**) The fish’s electric organ discharge (EOD) frequency shifts up continuously as the jamming signal is being applied. The area shaded with light blue indicates when jamming is turned on and lasts 40 seconds. The jamming signal frequency (current amplitude = 25 mA) is set to be 5 Hz below the fish’s EOD frequency. The fish’s EOD frequency gradually declines back to its normal value after jamming is turned off after around 60 seconds. (**B**) Maximum frequency shift during the fixed 40 second jamming window as a function of jamming current amplitude. Maximum frequency shift is averaged for every jamming amplitude across a total of 26 trials (*n* = 6 for 0 mA, *n* = 5 for 5 mA, *n* = 3 for 10 mA, *n* = 5 for 25 mA, *n* = 7 for 50 mA). The amount of jamming applied has a clear positive correlation with the EOD frequency shift.

**Figure 4—figure supplement 1.**
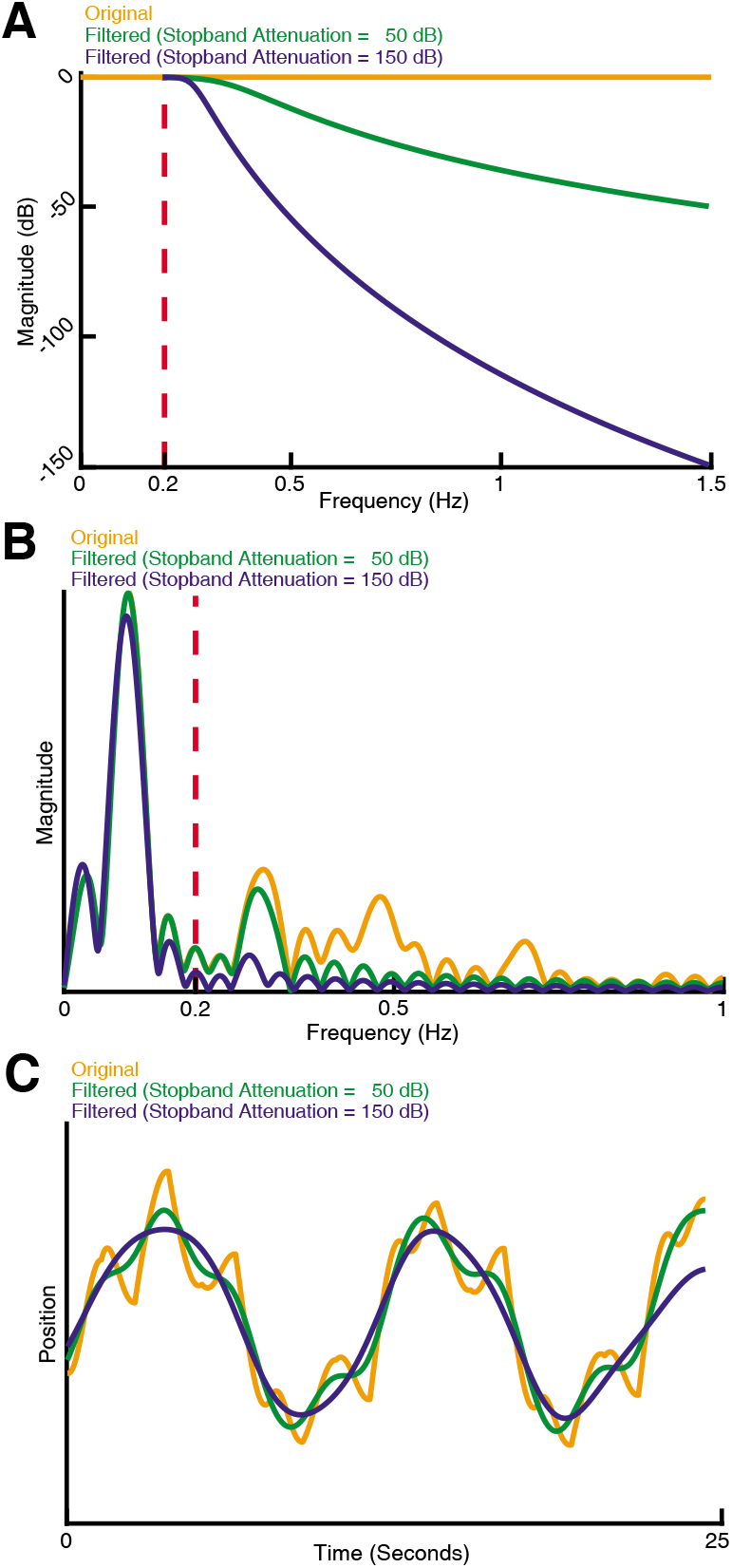
How sensing-related movements were attenuated for analyzing the impact of their diminishment. (**A**) Response of three types of sensing-related movement attenuation filters. We used an IIR lowpass filter with a cutoff frequency of 0.2 Hz to avoid filtering the baseline tracking frequency band of the target (0.1 Hz in this case). In the two filtered examples shown, the attenuation at 1.5 Hz is set to 50 dB and 150 dB to achieve progressively stronger attenuation of sensing-related motion. (**B**) Frequency response of the original and filtered trajectory. The baseline tracking gain between 0 Hz to 0.2 Hz is well preserved while the sensing-related motion frequency band between 0.2 Hz to 1.5 Hz is attenuated according to the attenuation gain. (**C**) Comparison of original and filtered trajectories. As the stopband attenuation increases, the amplitude of sensing-related movements decrease.

**Figure 6—figure supplement 1.**
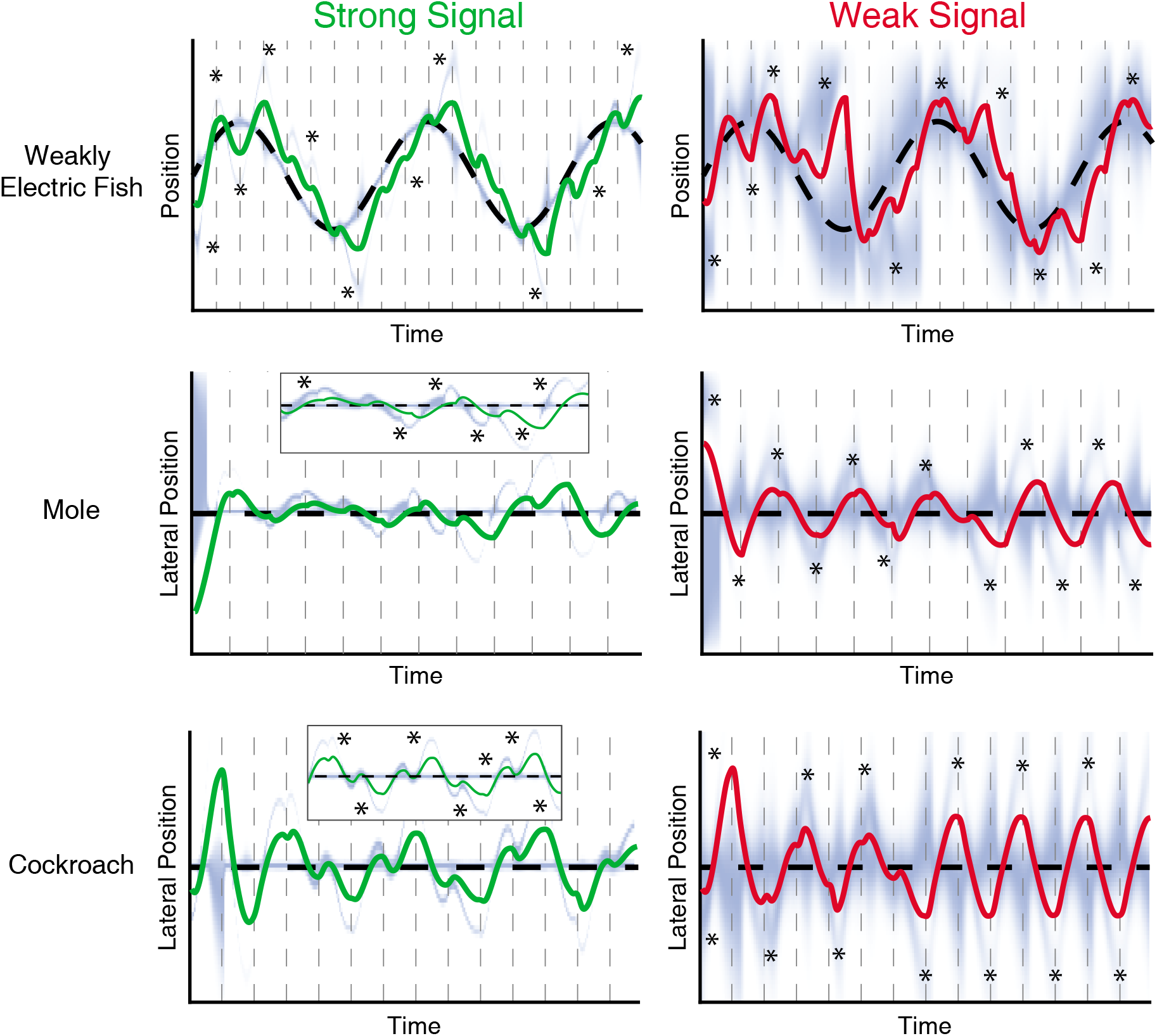
Evolution of belief over time for the trials shown in Figure 6. EIH simulations of tracking behavior of weakly electric fish, mole, and cockroach. The trials shown are identical to those shown in Figure 6. Simulated sensor position over time is the solid green line for the strong signal condition, and the solid red line in the weak signal condition. The target position is the dashed black line. The repeated dashed gray lines mark each planning update event that segment the trajectory into individual planned trajectory segments of length *T*. The blue heatmap shows the progression of the belief over time. Note the presence of fictive distractors, where the belief distribution becomes multi-peaked (asterisks). The inset in the mole and cockroach trials show a magnified segment to better visualize these distractors in the strong signal case.

**Figure 6—figure supplement 2.**
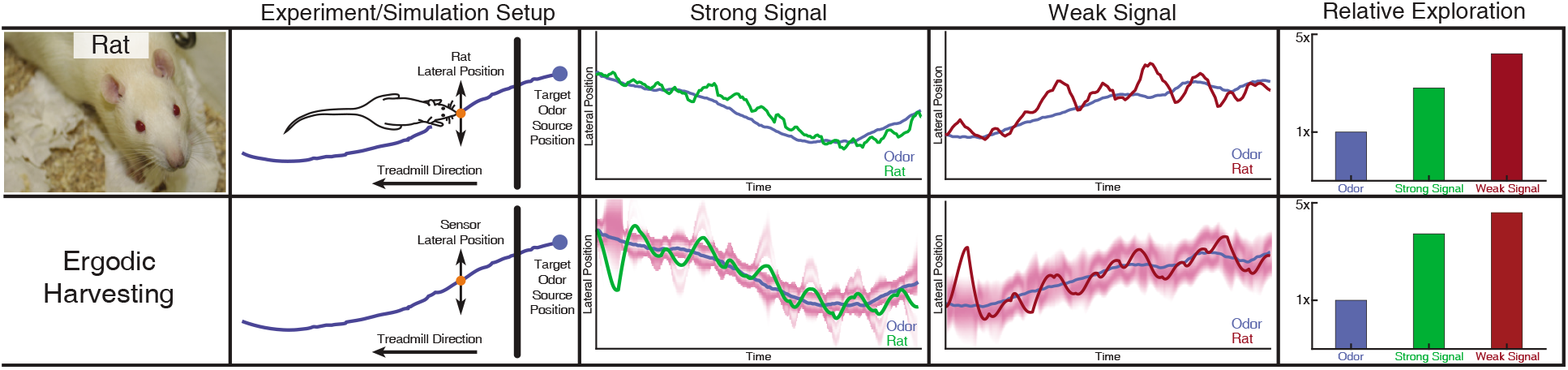
Rat odor tracking behavior and EIH simulation. In a prior study (***Khan et al., 2012***), Wistar rats (*Rattus norvegicus,* Berkenhout 1769) performed an odor tracking task by following a uniform odor trail on a moving treadmill with only olfactory information, under two different surgical conditions: with one side of the nose blocked (single-side nares stitching) and no blocking (sham stitching) as a control group. The experiment setup and tracked trajectory is shown in the top panel. Position data on the axis parallel to the treadmill were ignored, while positions on the perpendicular axis were recorded. The relative exploration was computed as with electric fish, using the cumulative 1D distance traveled by the rat divided by the cumulative distance of the transverse location of the odor trail. The olfactory system of the rat was modeled in the same fashion as for electric fish: as eventually providing an estimate of the location of the odor trail, modeled as a single 1D point-sensor of location with a Gaussian probability distribution controlled by the SNR of simulation (Methods). The blocked nares condition is here counted as the weak signal with lower SNR, and the sham stitching condition is counted as the strong signal. In the second row, we show EIH simulations of the weak and strong target trajectories, with the EID profile overlaid in magenta. It shows similar predicted sensor trajectories with a similar change in the relative exploration between these signal conditions as measured. These data are not included in the main study due to the low number of trials. Photo credit: original photo by Jean-Etienne Minh-Duy Poirrier, courtesy of Wikipedia, Creative Commons Attribution-Share Alike 2.0 Generic license.

**Figure 6—figure supplement 3.**
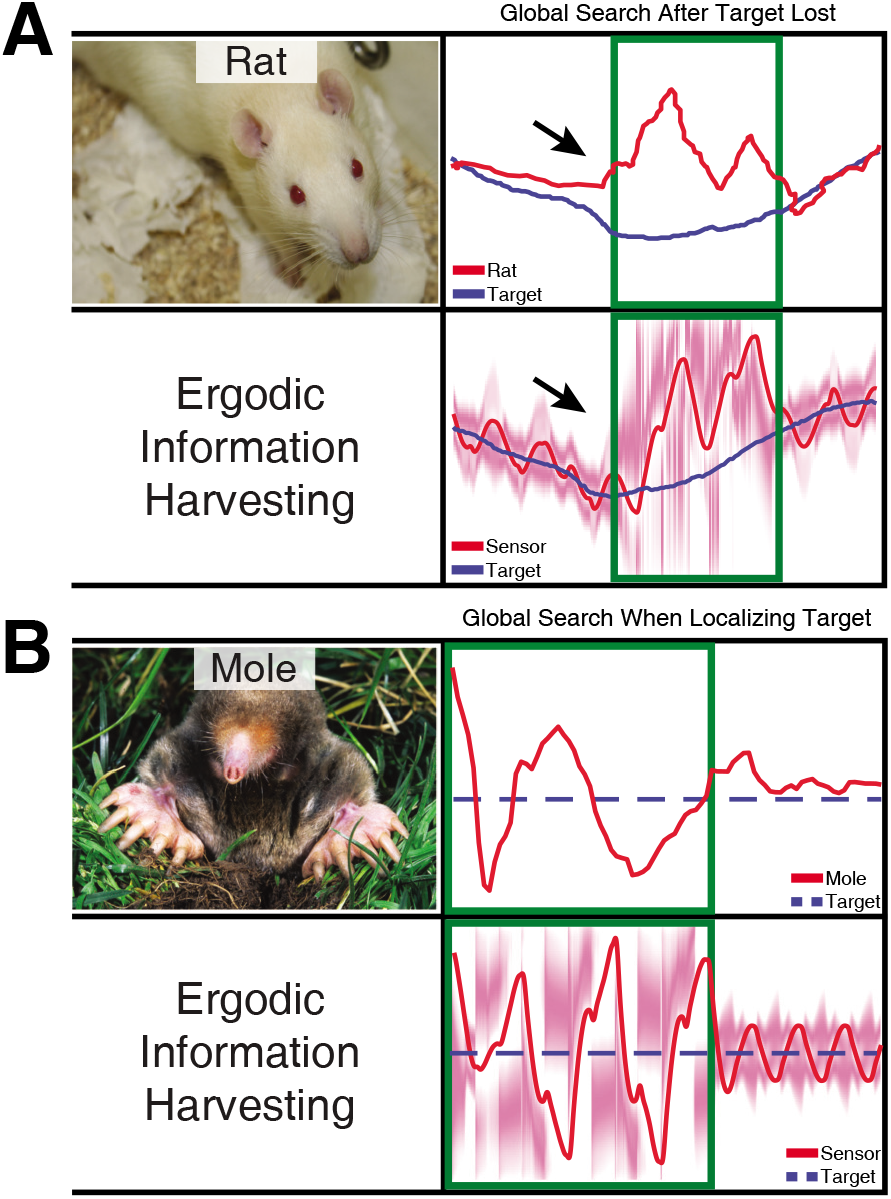
Measurements compared to prediction of EIH in two conditions where an animal needs to find the signal during tracking behavior. As a demonstration of how our model seamlessly transitions between exploitation and exploration, we examined an instance in the measured behavior of the live animals in which the rat appears to lose a signal it is following (***Khan et al., 2012***), and an instance in which the mole starts off without finding the path to the source (***Catania, 2013***). The measured result shown here for both animals was wider casting motions transverse to the direction of movement. We examined whether these two behaviors (wider casting with loss of signal during tracking or when not finding the signal initially) would be predicted by EIH. (**A**) To simulate the condition of losing a signal trail in the middle of a successful tracking trial, we ran EIH as for the rat olfactory tracking case of Figure 6—figure supplement 2, but with the trajectory shown in in panel A. In the “target lost” region (green square in figure), the sensor’s input was replaced by random draws from the observation model. (**B**) To simulate the condition of starting without knowing where the trail is for an extended period of time, we ran EIH as for the mole olfactory tracking case of Figure 6B, but with the trajectory shown in panel B. In the “target not yet acquired” region (green square in figure), the sensor’s input was replaced by random draws from the observation model. In both instances, we observed wider casting motions transverse to the direction of travel similar to those measured (output from EIH below experimental data, overlaid with the EID profile in magenta). Similar to the measured behavior, our algorithm seamlessly transitions from exploitation to exploration when the expected information density becomes more diffuse. Photo credit: Rat (Original photo by Jean-Etienne Minh-Duy Poirrier, courtesy of Wikipedia, Creative Commons Attribution-Share Alike 2.0 Generic license);Mole (Original photo by Kenneth Catania, Vanderbilt University, courtesy of Wikipedia, Creative Commons Attribution-Share Alike 3.0 Unported license).

**Figure 6—figure supplement 4.**
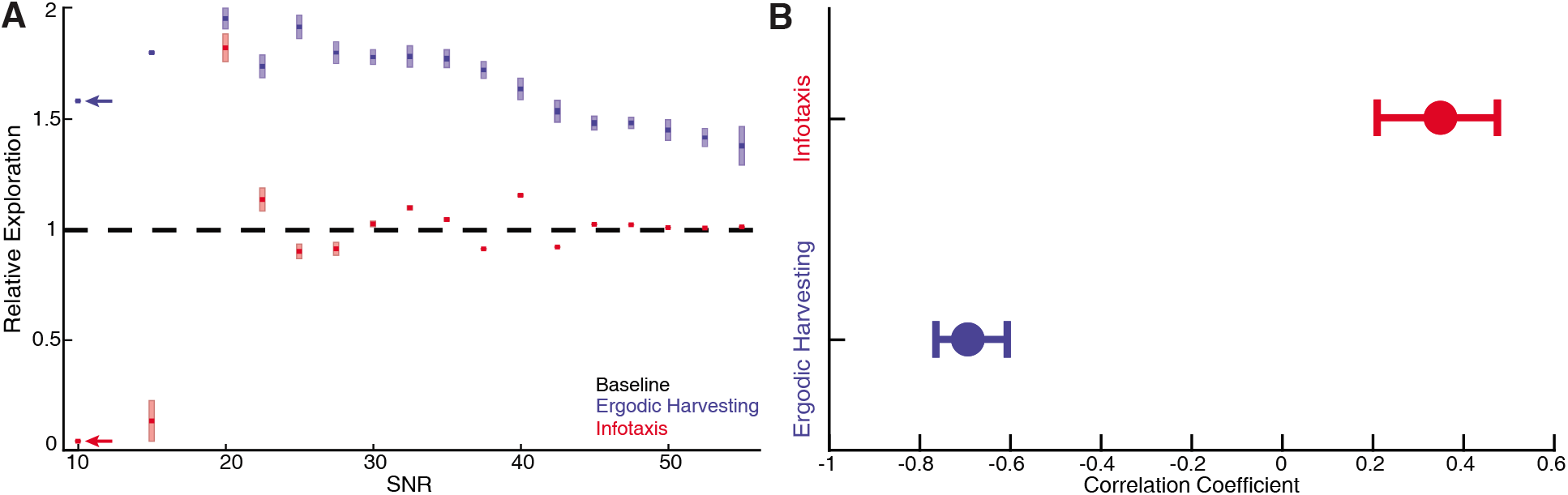
Systematic comparison between EIH and infotaxis in tracking a sinusoidally moving target. Simulations of a sensor tracking a target moving sinusoidally under 17 different SNR conditions from 10 dB to 55 dB. For each SNR condition, 10 simulations with different pseudo-number seeds are performed to establish the confidence interval (*n* = 170 for EIH, *n* = 170 for infotaxis). (**A**) As the SNR decreases from 55 dB, EIH exhibits elevated relative exploration (see Methods) in the form of increased sensing-related movements. This is consistent with the animal behaviors summarized in Figures 1–7. EIH’s elevated exploration with decreasing SNR tapers at ≈20 dB as the Bayesian filter fails to converge the posterior due to significantly less informative measurements. Infotaxis does not show the trend of increased exploration as signal weakens between 55-20 dB, and prescribed cessation of movement at the lowest (10 dB) signal strength. Throughout most of the frequency spectrum, infotaxis attempts to hug one edge of the EID, leading to near unity relative exploration. (**B**) The correlation coefficient between relative exploration and signal strength computed from **A** for EIH and infotaxis (*n*=170 trials for EIH, and *n*=170 for infotaxis). The vertical line indicates the 95% confidence interval. Ergodic harvesting has a significantly more negative correlation, indicating exploration increases as signal strength declines. The same trend is absent in infotaxis.

**Figure 6—figure supplement 5.**
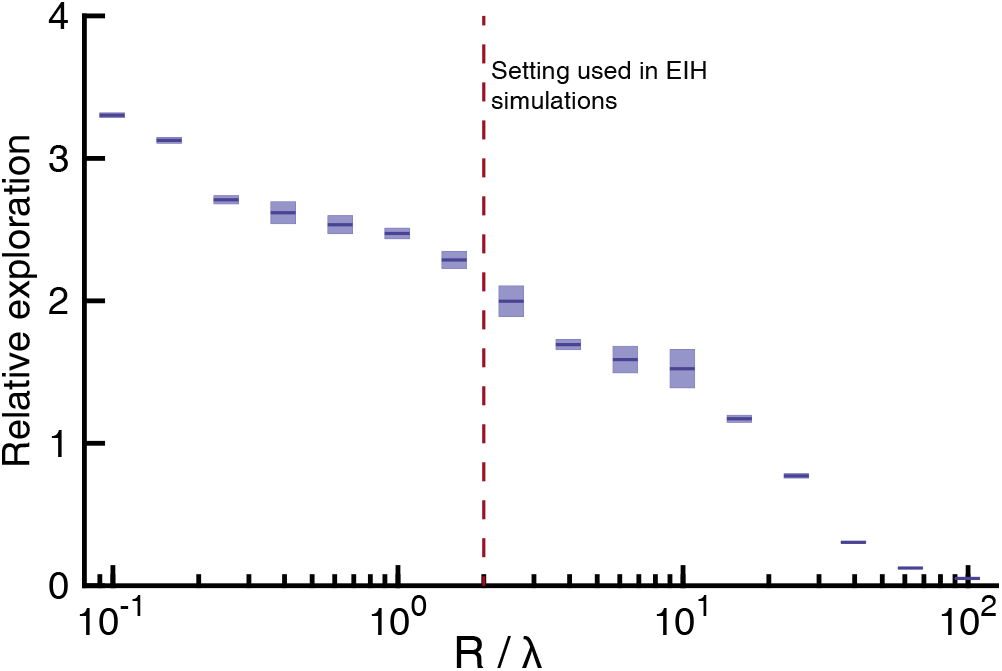
Sensitivity analysis on the ratio between control cost and ergodic cost in the objective function of trajectory optimization. Simulations are conducted in the same way as Figure 6—figure supplement 4 but only for a fixed SNR under weak signal conditions (20 dB). The control cost term *R* is varied while fixing the ergodic cost term *λ* to be 5 for all the simulations. We simulated 16 different ratios of *R/λ* (shown along x axis on a log scale), each done with 10 different pseudo-random number seeds to establish confidence intervals. The relative exploration of the simulated trajectories are shown in the y axis on a linear scale with the sample mean marked by the solid color line and 95% confidence interval marked by the vertical color patch. The *R/λ* ratio used for all the EIH simulations included in the results is shown by the red dashed vertical line. Overall, as the *R/λ* drops, relative exploration drops monotonically because movement incurs higher cost. The trend also suggests that as the *R/λ* ratio increases, movement will cease. The value of *R/λ = 2* that we used in our study results in a relative exploration value of around 2. The variation in relative exploration with an order of magnitude change in *R/λ* from 1 to 10 is approximately 2.5 to 1.5. The same sensitivity analysis was also done for a strong signal condition but is not shown as the trend is similar.

## List of Figures

- Figure 1 — Illustration of the information maximization and energy-constrained proportional betting strategies Figure 1—figure supplement 1 — Whole-body or sensory organ small-amplitude motions are ubiquitous as animals track targets
- Figure 2 — Longitudinal refuge tracking behavior in weakly electric fish and illustration of the Ergodic Information Harvesting hypothesis Figure 2—figure supplement 1 — Dual target tracking simulation with EIH and infotaxis Figure 2—figure supplement 2 — Single target tracking simulation with EIH and infotaxis in the presence of a simulated physical distractor Figure 2—figure supplement 3 — Effect of jamming and how it varies with jamming intensity
- Figure 3 — Systematic comparison between fish behavior tracking data and EIH predictions
- Figure 4 — Sensing-related movements reduce tracking error Figure 4—figure supplement 1 — How sensing-related movements were attenuated for analyzing the impact of their diminishment
- Figure 5 — Relative energy for electric fish tracking behavior and EIH-generated behavior with attenuated body oscillations
- Figure 6 — Trajectory comparison of animals tracking a target compared to EIH, and relative exploration across all trials Figure 6—figure supplement 1 — Evolution of belief over time for the trials shown in Figure 6 Figure 6—figure supplement 2 — Rat odor tracking behavior and EIH simulation Figure 6—figure supplement 3 — Measurements compared to prediction of EIH in two conditions where an animal needs to find the signal during tracking behavior Figure 6—figure supplement 4 — Systematic comparison between EIH and infotaxis in tracking a sinusoidally moving target Figure 6—figure supplement 5 — Sensitivity analysis on the ratio between control cost and ergodic cost in the objective function of trajectory optimization
- Figure 7 — Spectral analysis of live animal behavior and simulated behavior

## List of Tables

- Table 1 — Parameters of EIH Simulation
- Table 2 — Simulation Parameters Used For Each Figure

## List of Videos

- Video 1 —Overview of energy-constrained proportional betting theory.
- Video 2 — The ergodic information harvesting algorithm applied to control a bio-inspired robot.

## Notes

### Competing Interest Statement

The authors have declared no competing interest.

