## Appendices for "Tuning movement for sensing in an uncertain world"

List of Appendices


- Appendix 1 — Background on ergodicity
- Appendix 2 — Relationship between Ergodicity and Kullback-Leibler divergence
- Appendix 3 — How the distance from ergodicity is computed
- Appendix 4 — How the expected information density (EID) is defined and computed
- Appendix 5 — Comparison to stochastic models
- Appendix 6 — EIH in 2-D and 3-D behavioral contexts


### Appendix 1

#### Background on ergodicity

Ergodicity plays an important role in multiple scientific disciplines particularly in stochastic systems and statistical mechanics. In the setting of Markov chains, defined by states and stochastically-driven transitions between states, a system is ergodic if every aperiodic path that leaves a given state must return to that state with probability one (***Lasota and Mackey, 2013***). However, in the present work we are not interested in stochastic evolutions—though there is the possibility that stochasticity in a system could contribute to coverage needs, something discussed momentarily. Instead, we are interested in deterministic decisions—*i.e.*, control decisions—that provide coverage with respect to regions of high information density.

A key insight from ***Mathew and Mezic*** (***2011***) was to use the definition of ergodicity for dynamical systems—that a trajectory $x(t)$ spends time in any particular neighborhood $\mathcal{N}$ proportional to the measure of the distribution $\Phi$ over that neighborhood $\int_{\mathcal{N}} \Phi(x)dx$—to create a metric on deterministic trajectories. That is, once $\Phi$ is given, there is nothing stochastic in the question of coverage. Instead, there is only the question of how much coverage a particular trajectory $x(t)$ provides relative to $\Phi$. It should additionally be noted that coming up with a metric is not trivial, in large part because any mathematical comparison must be able to compare two distinct mathematical ideas—a distribution and a trajectory. A distribution is a probability over a region, while a trajectory is a continuum of states parameterized by time $t$. In ***Mathew and Mezic*** (***2011***) the authors note two critical steps in creating such a comparison. First, they note that a trajectory can *also* be represented as a sequence of Dirac delta functions $\delta(x)$ also parameterized by time $t$, so that the comparison is between two distributions rather than between a distribution and a trajectory. Secondly, perhaps more importantly, they use the fact that spatial Fourier transforms are well posed for quite general distributions, including Dirac delta functions. From these two steps they conclude that the coefficients of the Fourier transform provide an infinite set of variables that can be used to form a metric. Importantly, none of this analysis requires the trajectories to be stochastic, though similar analysis can be done for stochastic executions (where, for instance, one can imagine that at least small amounts of noise might improve coverage locally).

### Appendix 2

#### Relationship between Ergodicity and Kullback-Leibler divergence

Ergodicity provides a mathematical approach for comparing a trajectory to a distribution in a way that is similar to how K-L divergence compares a distribution $P$ to a distribution $Q$. However, K-L divergence cannot be directly applied to trajectories. This is because an idealized trajectory (no uncertainty by itself) is an aggregations of singletons in the form of individual Dirac delta functions (each of zero variance), one for each time $t$. This leads to infinite K-L divergence because the K-L divergence measures how much information changes when using one distribution to represent another distribution (this is why the K-L divergence is often called the relative entropy). In the case of representing a distribution with a delta function, infinite information has been gained because the argument of the delta function is specified with zero variance. As a concrete special case, if one approximates a delta function with normal distributions of decreasing variance, the entropy goes to infinity as the variance goes to zero.

Here is a brief illustration of some of the steps needed to show why K-L divergence will not work for trajectories for those who would like more explanation. This illustration elides a number of technical issues that would need to be carefully worked through for a more rigorous result. Imagine we have a 1-D trajectory that consists of two points (*i.e.* singletons): $\{0, 1\}$. We can call this a probability density function (PDF) $P$ consisting of two Dirac deltas, $\delta(x-0)$ and $\delta(x-1)$—that is, $P(x) = \frac{1}{2}(\delta(x-0) + \delta(x-1))$. The differential probabilities $P(0) = P(1)$ both integrate to $\frac{1}{2}$ since the total probability is $1$. Suppose we want to compute the K-L divergence between $P$ and $Q$, where $Q$ is an arbitrary Gaussian distribution with a mean of 0.5 and a non-zero variance. According to the general form of computing K-L divergence:

$$\begin{aligned} D_{KL}(P||Q) &= \int_x P(x) \log \frac{P(x)}{Q(x)} dx \\ &= \int_x P(x) \log P(x) dx - \int_x P(x) \log Q(x) dx \end{aligned} \tag{1}$$

where $\int_x P(x) \log P(x) dx$ is the differential entropy of $P$, which is undefined in this case. To understand why, consider an arbitrary Gaussian distribution, $f(x) \sim \mathcal{N}(\mu, \sigma)$. Computing the first term of the expression for differential entropy (Eq. 1) gives $h(f) = \int_x f(x) \log f(x) dx = \log(\sigma) + \frac{1}{2}\log(2\pi e)$, which is undefined in the limiting case of a Dirac delta function with $\sigma = 0$ since $\lim_{\sigma \to 0^+} \log(\sigma) = -\infty$. Hence, the K-L divergence between a Dirac delta function (representing the idealized trajectory) and a smooth (EID) distribution is undefined. (Note that the other term in the K-L divergence that depends on $Q(x)$ will evaluate to a constant, so does not impact the well-posedness of K-L divergence for a trajectory.)

Similar to K-L divergence, mutual information, which quantifies the amount of information obtained about one random variable $X$ by observing another random variable $Y$, defined as $I(X;Y) = D_{\text{KL}}\left(P(X,Y) \;||\; P_X P_Y\right)$, is another widely used approach for quantifying information between two distributions of random variables. For jointly discrete or jointly continuous pairs $(X, Y)$, it is the K-L divergence between the joint distribution $P(X,Y)$ and the product of the marginal distributions $P_X$ and $P_Y$. Given that the K-L divergence between a trajectory and a distribution is undefined as discussed above, mutual information also cannot be applied to trajectories. More generally, as we stated in the main text that animal physical trajectories are considered the behavior of dynamical systems rather than sample paths from a stochastic process, methods like K-L divergence and mutual information that requires both inputs to be distributions are undefined and hence will not work in the case where one of these is a trajectory.

#### Appendix 3

##### How the distance from ergodicity is computed

These details are largely from our prior publication ***Miller et al.*** (***2016***), repeated here for convenience. The spatial statistics of a trajectory $x(t)$ are quantified by the percentage of time spent in each region of the workspace

$$C(x) = \frac{1}{T} \int_0^T \delta[x - x(t)] dt \tag{2}$$

where $\delta$ is the Dirac delta function and $T$ is the duration of the trajectory. The distance from ergodicity is then defined as the sum of weighted squared distances between the Fourier coefficients of the EID $\phi_k$ and the coefficients $c_k$ of the distribution representing the time-averaged trajectory. The Fourier coefficients $\boldsymbol{\phi_k}$ of the distribution $\Phi(\boldsymbol{x})$ are computed using an inner product

$$\phi_{\boldsymbol{k}} = \int_X \phi(\boldsymbol{x}) F_{\boldsymbol{k}}(\boldsymbol{x}) d\boldsymbol{x} \tag{3}$$

and the Fourier coefficients of the basis functions along a trajectory $\boldsymbol{x}(t)$, averaged over time, are calculated as

$$c_{\boldsymbol{k}}(\boldsymbol{x}(t)) = \frac{1}{T}\int_0^T F_{\boldsymbol{k}}(\boldsymbol{x}(t))dt \tag{4}$$

where $T$ is the final time and $F_k$ is a Fourier basis function that takes the form of

$$F_k(x) = \frac{1}{h_k}\prod_{i=1}^{n}\cos\left(\frac{k_i\pi}{L_i}x_i\right) \tag{5}$$

where $h_k$ is a normalization factor ***Mathew and Mezic*** (***2011***) and $L_i$ is a measure of the length of the dimension. Finally, the ergodic metric is specified as

$$\mathcal{E}(x(t)) = \sum_{k=0}^{\mathbf{K}} \Lambda_k [c_k(x(t)) - \phi_k]^2 \tag{6}$$

where $\mathbf{K}$ is the number of Fourier coefficients used along every one of the $n$ dimensions and $\Lambda_k = (1+\|k\|^2)^{-s}$, where $\Lambda_k$ come from the Sobolev space norm and $s = \frac{n+1}{2}$ places more weight on lower frequency information ***Mathew and Mezic*** (***2011***). Given the definition above, the ergodic metric $\mathcal{E}(x(t))$ quantifies the difference between a given distribution of EID and the spatial statistics $C(x)$ of an trajectory $x$. We say a trajectory is perfectly ergodic with respect to an EID map if $\mathcal{E}(x) = 0$, that is, the spatial statistics of $x$ exactly matches the distribution of the target EID map.

#### Appendix 4

##### How the expected information density (EID) is defined and computed

Given an unknown random variable $\boldsymbol{\theta}$ to estimate (in the context of tracking simulations, $\boldsymbol{\theta}$ represents the location of the tracking target), EIH evaluates an expected information density EID$(x)$ at every planning update based on the current belief $p(\boldsymbol{\theta})$. The EID essentially answers the following question: given the probability of $\boldsymbol{\theta}$ being a particular value, and given the likelihood of receiving a particular voltage $V$ corresponding to that value, what is the average amount of information we expect to receive by visiting a state $x$?

Computing EID$(x)$ requires several steps. First, we define a Gaussian likelihood function that predicts how likely the sensor is to obtain a measurement $V \in \mathcal{V}$ given the current belief $p(\boldsymbol{\theta})$, where $\mathcal{V}$ is the set of all possible sensor measurements (see Chapter 7.2 of ***Robinson 2016*** for details regarding the likelihood function).

$$p(V \mid \boldsymbol{\theta}, x) = \frac{1}{\sqrt{2\pi}\sigma}\exp\left[-\frac{(V - \Upsilon(\boldsymbol{\theta}, x))^2}{2\sigma^2}\right] \tag{7}$$

Here $x \in \mathcal{X}$ is the location of the sensor ($\mathcal{X}$ is the space of all possible sensor locations), and $\Upsilon(\boldsymbol{\theta}, x)$ is the observation model assuming a known target location $\boldsymbol{\theta}$ evaluated at sensor location $x$.

Next, with a predicted distribution of measurements for each choice of $x$ from the likelihood function $p(V|\boldsymbol{\theta}, x)$, we evaluate what the new posterior belief $p(\boldsymbol{\theta} \mid V, x)$ is expected to be if the sensor were to take a hypothetical measurement at a given location $x$ in the workspace. From the multiplication rule of conditional probability (see Eq. 3.16 of ***Kokoska and Zwillinger 2000***),

$$P(A \cap B) = P(A|B)\ P(B) = P(B|A)\ P(A) \tag{8}$$

where $A$ and $B$ are two random events, we obtain the Bayes update rule:

$$P(A|B) = \frac{P(B|A)\ P(A)}{P(B)}. \tag{9}$$

For each choice of potential $x$ where a sensor measurement could be taken, the new posterior is therefore computed by (see Chapter 3.3.9 of ***Kokoska and Zwillinger 2000*** and Chapter 2.4 of ***Thrun et al. 2005***):

$$\begin{aligned} p(\boldsymbol{\theta} \mid V, x) &= \eta\ p(\boldsymbol{\theta} \mid x)\ p(V \mid \boldsymbol{\theta}, x) \\ &= \eta\ p(\boldsymbol{\theta})\ p(V \mid \boldsymbol{\theta}, x) \end{aligned} \tag{10}$$

where $\theta$ corresponds to $A$ and $V$ corresponds to $B$ in Eq. 8. In Eq. 10, $p(\theta \mid x) = p(\theta)$ because $\theta$ and $x$ are mutually independent, and $\eta = \frac{1}{p(V \mid x)} = \frac{1}{\int_{\theta} p(V \mid \theta, x)\; p(\theta) d\theta}$ is a normalization factor that constrains the posterior belief $p(\theta \mid V, x)$ to be a probability distribution (see Chapter 2.4 of ***Thrun et al.*** (***2005***)).

Given a posterior belief $p(\theta \mid V, x)$ evaluated on a potential $V$ measured at a potential location $x$, the entropy reduction from the prior belief $p(\theta)$ can be evaluated using:

$$\Delta S(V, x) = S\left[p(\theta)\right] - S\left[p(\theta \mid V, x)\right] \tag{11}$$

where $S\left[p(\theta)\right] \in \mathbb{R}^1$ and $S\left[p(\theta)\right] = -\sum_{\theta} p(\theta)\; \log p(\theta)$ is the Shannon-Weaver entropy of the prior belief $p(\theta)$, while $S\left[p(\theta \mid V, x)\right]$ is the Shannon-Weaver entropy of the posterior belief.

For any given prior belief $p(\theta)$, the probability of the sensor receiving a measurement $V$ given a choice of sensing location $x$ is not necessarily constant. Therefore, to evaluate the expected entropy reduction at a given sensing location $x$, the entropy reduction $\Delta S(V, x)$ needs to be weighted by the measurement probability $p(V \mid x)$ that is consistent with the prior belief $p(\theta)$. This weighted probability can be obtained by applying the law of total probability (see Eq. 3.17 of ***Kokoska and Zwillinger 2000***) to the normalized likelihood function $p(V \mid \theta, x)$ treated as a probability distribution (see Chapter 5.2 of ***Robinson 2016***).

$$p(V \mid x) = \int_{\theta} p(\theta) p(V \mid \theta, x) d\theta \tag{12}$$

Finally, the expected information density at location $x$—EID$(x)$—is obtained by computing the mathematical expectation (see Chapter 3.5.1 of ***Kokoska and Zwillinger 2000***) of the entropy reduction *if one were* to take a measurement at location $x$. That is, EID$(x)$ is the weighted average entropy reduction resulting from the conditional probability $p(\theta \mid V, x)$, weighted by the measurement probability $p(V \mid x)$.

$$\text{EID}(x) = \text{E}\left[\Delta S(x)\right] = \int_{V} p(V \mid x)\; \Delta S(V, x) dV \tag{13}$$

An interactive online tutorial and video describing the above steps of EIH is available (***Chen et al., 2020***).

#### Appendix 5

##### Comparison to stochastic models

If animal trajectories are sample paths of a process made up of deterministic and stochastic parts, then some observed small-amplitude oscillations can be modeled by a stochastic search process, similar to that reported in *Drosophila* (***Reynolds and Frye, 2007***; ***Censi et al., 2013***; ***Mongeau and Frye, 2017***; ***Ferris et al., 2018***). Since a sensed target is always present in the tracking behavior analyzed here, it is unlikely that the trajectories analyzed here are purely driven by stochastic search with no intended target (*e.g.* refuge, food source, odor plume). Most models that implement stochasticity by drawing actions based on the EID distribution represent stochasticity in an abstraction of the space in which the body evolves based on its physics (e.g., a Thompson sampling process that randomly samples locations, ignoring the physics and energetics of getting to those locations). If a stochastic signal is directly driving the physics of the body, small random walks will indeed occur, but large-scale motion of the entire body will not occur unless the physical randomness is very large. Moreover, stochastic search can be considered a special case of ergodic search. In general, a random walk will lead to coverage of some area, and that same area could be covered using the ergodic coverage algorithms described here. But the ergodic coverage algorithms enable an animal to adapt as the environment changes, where a given stochastic search will be independent of changes in the environment. This difference may matter in settings where changes in environment matter to search success. Such a scenario is demonstrated in Figure 6—figure supplement 3 where we provide two examples of target loss. In both cases, the animal exhibits immediate local-to-global search transitions which is naturally reproduced by EIH. Finally, the ergodic harvesting strategy can be applied to stochastic scenarios, where the dynamics include a stochastic process, as shown in ***De La Torre et al.*** (***2016***) without substantially impacting the solution.

#### Appendix 6

##### EIH in 2-D and 3-D behavioral contexts

The presented animal behavior analysis and EIH simulations are limited to a 1-D workspace in the presence of a single target moving along a line. While all animals move in 3-D, the behaviors we examined were minimally distorted by projection to 1-D. Here we consider extensions to behaviors that cannot be projected to 1-D.

Although we show evidence suggesting that EIH naturally balances the exploration-versus-exploitation trade-off in the case of signal loss in 1-D (see Figure 6—figure supplement 3), it is unclear how EIH would behave in similar cases in workspaces with more than one dimension. For example, ***Calhoun et al.*** (***2014***) shows that infotaxis in a 2-D context will respond to a local distractor in the EID by first going straight towards the peak, as also shown in Figure 1B, and then engage in circular motion around the distractor peak as it gradually gets rejected by new observations. As a comparison, in Figure 1C we show that EIH predicts that the animal will not wander around the distractor peak for long, but rather dwell in such distractor peaks to make additional observations before naturally switching to other regions including the true target location. Such behavior emerges from EIH without any dependence on changes in the EID, whereas infotaxis is dependent on changes in the EID. This also applies to Figure 6—figure supplement 4 at 20 dB SNR, one of few places where infotaxis exhibits a relative exploration level similar to EIH. This increase in sensor movement is driven by changes in the information landscape due to a very high level of uncertainty. However, the movements of infotaxis at this SNR does not generate systematic coverage with respect to the EID and actually leads to sub-optimal tracking performance ($64 \pm 3$% tracking error for EIH versus $74 \pm 4$% tracking error for infotaxis at 20 dB; Kruskal-Wallis test, $p < 0.001$, $n = 20$). Further investigation is needed to explore the effect of temporal variation in the information landscape on sensing behavior—in 2-D contexts such as explored by ***Calhoun et al. 2014*** and in 3-D contexts—for insight into on how animals approach the exploration-versus-exploitation trade-off in various scenarios such as signal loss.
